# Perturbation-based mapping of natural frequencies with direct intracranial stimulation of the human brain

**DOI:** 10.1101/718064

**Authors:** Julià L Amengual, Chloé Stengel, Tristan Moreau, Claude Adam, Mario Chavez, Antoni Valero-Cabré

## Abstract

Theoretical and experimental evidence suggest that the induction of oscillatory activity by an external rhythmic source on a specific brain area is maximally efficient if the input pattern matches its so-called ‘natural’ frequency, defined as the predominant neural rhythm at which the activity of this area tends to fluctuate spontaneously. Based on this principle, single pulse Transcranial Magnetic Stimulation (TMS) coupled to scalp electroencephalography (EEG) has provided evidence of frequency-specific power increases within a unique ‘natural’ frequency band, considered common to the whole lobe.

In an attempt to gain deeper insight into this phenomenon and set the basis for a finer-grained atlas of ‘natural’ frequencies, here we analyzed intracranial EEG (iEEG) signals modulated by single pulses of direct electrical brain stimulation in human patients implanted with depth multielectrodes. Our analyses revealed changes in local EEG activity emerging from local oscillators and contributing to a complex distribution of frequency-specific ‘natural’ rhythmic responses throughout cortical regions. Moreover, challenging the notion of ‘natural’ oscillations featuring a predominant frequency band characteristic for an entire lobe, our data support a rich diversity of spectral fingerprints (narrowband, vs. broadband or multiband) with single or multiple frequency peaks, often encompassing contiguous frequency bands, operating at a very local scale.

Our findings contribute novel insights on which specific brain areas could be more likely to be synchronized at a given frequencies band and their preferred coupling frequencies, features that could ultimately inform on their structural and functional organization. Our results may also increase our mechanistic understanding of invasive and noninvasive brain stimulation and promote further developments of these approaches for the manipulation of brain oscillations subtending normal and impaired cognition.

## INTRODUCTION

Cutting-edge research conducted during the last decade suggests that oscillatory signals can be shaped by task-dependent synchronization or desynchronization of brain rhythms enabled locally and spreading throughout distributed networks^1, 2^. In the behavioral domain, brain oscillations have been found to play an instrumental role in coding for specific aspects of human cognition such as the orienting of spatial attention, perceptual modulations, memory acquisition/consolidation, motor planning and action control^3–6^.

Lately, the use of non-invasive brain stimulation technologies such as Transcranial Magnetic Stimulation (TMS) or Transcranial alternating current stimulation (tACS) to explore the anatomical basis, neurophysiological coding and cognitive contributions to brain synchrony^7–11^ has gained momentum. Moreover, similar principles are also being applied to the development of more efficient non-invasive brain stimulation approaches aiming to modulate the symptoms of neuropsychiatric diseases^12^.

Indeed, recent reports have shown that frequency-tailored entrainment of oscillatory activity can be achieved either with isolated pulsed perturbations inducing local phase-resetting and the synchronization of local oscillators^13^, or alternatively, with periodical perturbations able to entrain oscillations at frequencies dictated by the temporal distribution of pulses within stimulation bursts^10, 14^. Moreover, combined (EEG)/Magnetoencephalography (MEG) recordings in humans have contributed neurophysiological evidence in favor of enhanced neural oscillatory activity during the delivery of focal patterns of rhythmic non-invasive (TMS/tACS) and invasive intracranial electrical stimulation^12, 14–16^.

Computational models developed to simulate oscillatory entrainment driven by external sources have suggested that the impact of rhythmic patterns could be maximized if the temporal distribution of the input matches the frequency of the to-be-entrained oscillators ^1^. Such approaches prone that oscillatory entrainment could be most easily achieved by tuning stimulation frequency to the most prominent rhythm operating physiologically on the targeted brain region, hence at the so called ‘natural’ frequency^13, 14^. However, mapping efforts exploring the precise distribution of ‘natural’ rhythms across brain systems and characterizing local ‘natural’ spectral fingerprints associated to specific brain sites remain a work in progress.

The most common strategy to compile atlases of ‘natural’ rhythmic patterns in the human brain has consisted in studying the spectral components of resting state neurophysiological signals with high resolution EEG/MEG systems ^17^. Nonetheless, the large intra-individual^18^ and inter-individual ^19^ variability of non-invasive electrophysiological recordings, and limitations shown by these methods to establish causal links between spectral fingerprints and discreet cerebral regions^13^ had limited the spatial resolution and reliability offered by such pioneering approaches. Additionally, the summed dynamics of multiple oscillators with random phase relations recorded extracranially using non-invasive methods tend to nullify the population sum, preventing also an accurate assessment of ‘natural’ frequencies of local neural populations at rest.

Pursuing a credible causal alternative, seminal work carried out in this domain by Rosanova et al. applied single pulses of Transcranial Magnetic stimulation (TMS) coupled to high-density scalp EEG and perturb spontaneous oscillatory dynamics in representative regions of the human cortex^13^. By measuring the power of the most prominent oscillations boosted by stimulation, this pioneering work identified the alpha, beta and beta-gamma as the ‘natural’ rhythms of occipital, temporal and frontal lobar regions, respectivele. Nonetheless, the limited ability of TMS to probe a large number of intra-lobar cortical sites (forcing a generalization of local outcomes across very large brain-regions) and standing uncertainties of non-invasive methods such as TMS (e.g. unclear mechanism of activation) and scalp EEG (e.g. source localization problem) call for alternative approaches.

Addressing this challenge, we here studied patterns of oscillatory activity induced by single pulses of intracranial electrical stimulation through intracranial EEG (iEEG) recordings in a cohort of epilepsy patients implanted with depth multi-electrodes ^17^. Datasets were acquired during invasive mapping sessions using single electrical pulses to causally localize and probe epileptogenic regions, prior to neurosurgical removal. The impact of electric stimulation was analyzed in the frequency domain by recording iEEG activity from multi-electrode contacts monitoring non-epileptogenic regions. The spectral signatures induced by electrical stimulation were integrated in a common human brain atlas presented in normalized MRI space.

We hypothesized that region-specific spectral fingerprints of varying complexity (e.g. unimodal vs. multi-modal frequency distributions) characterizing local ‘natural’ oscillatory activity would emerge in response to single pulse electrical perturbations and eventually outlast them briefly in a frequency dependent manner. We also predicted that the distribution of predominant ‘natural’ frequencies extracted from site-specific spectral fingerprints would prove rich and diverse, but respect the boundaries between lobar areas established previously with non-invasive approaches^13^. Collaterally, we expected physiological responses to single electrical pulses to follow state-dependent principles, hence be modulated by the phase of the ongoing local oscillations (at the ‘natural’ frequency) on the stimulated area at the time of the perturbation. Importantly, we speculated that fingerprint complexity could potentially inform on the local structural and physiological organization of the perturbed site and its ability to influence network dynamics.

## MATERIALS AND METHODS

### Patient populations and intracranial iEEG electrode montage

We studied intracranial EEG datasets from 19 medication-resistant epilepsy patients (11 females, 8 males) between 18 and 41 years old (mean age 24) implanted with intracranial deep electrodes (Figure 1) undergoing a thorough clinical evaluation of epileptic foci prior to consider their neurosurgical resection (Table 1). The study was sponsored by the *Institut Nationale de la Santé et la Recherche Médicale* (INSERM) and approved by the ethical committee (CPPRB, Comité Consultatif de Protection des Personnes participant à une Recherche Biomédicale) Ile-de-France I (reference number C11-16, 5-04). All methods were performed in accordance with the National (France), European (EU) and International (Declaration of Helsinki) guidelines and regulations.

**Figure 1.**
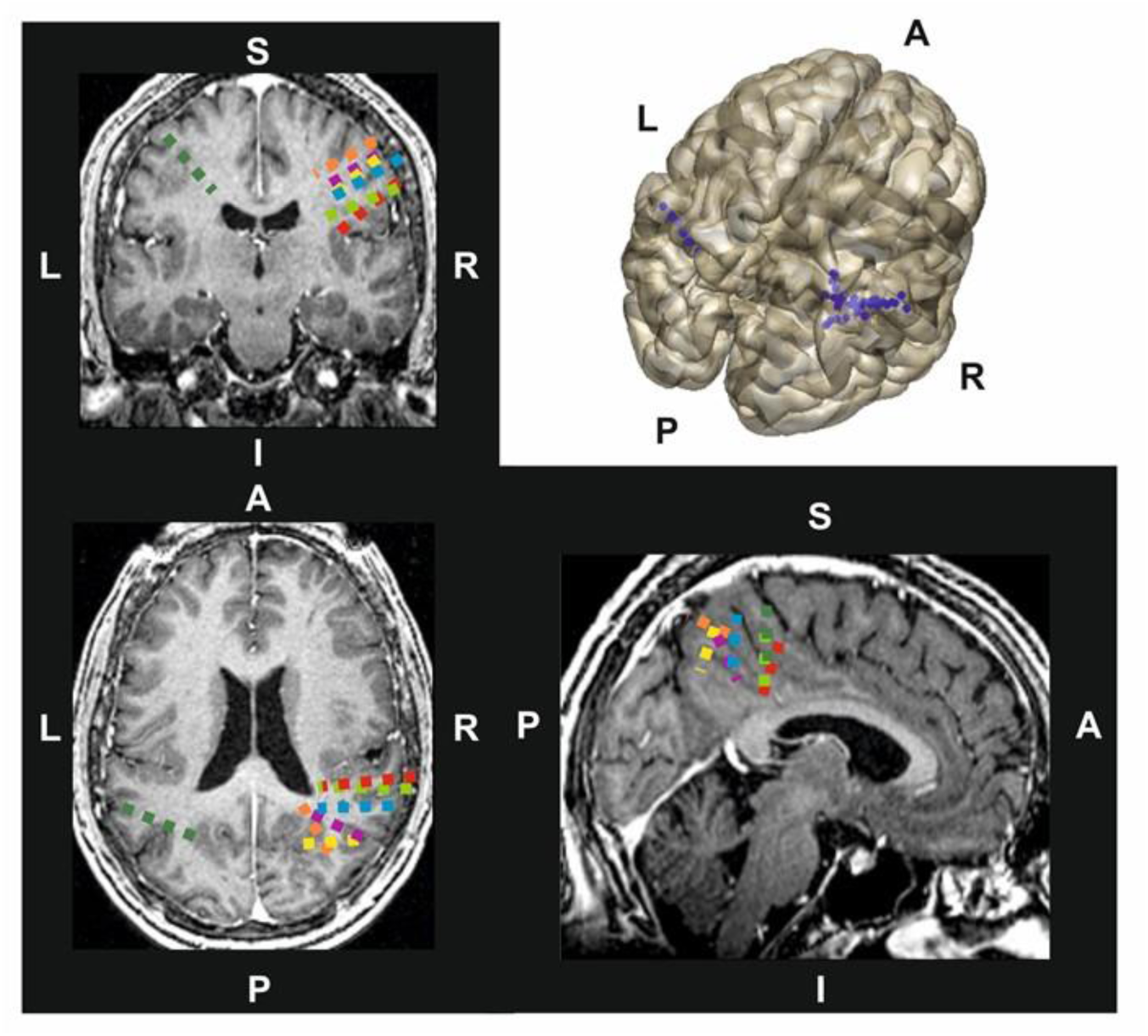
Individual multielectrode implantation schemes. Detailed view of the multielectrodes implantation scheme and contact localization on an T1-3D MRI volume (coronal, sagittal and axial views) from a representative patient of our dataset. Notice for this specific case multielectrode contacts are placed in sites within the right and left parietal lobe. Blue dots label the projection of each contact site onto the cortical surface of the brain

**Table 1.** Demographic data of each patient and implantation schemes. Table includes patient code, gender (**M**: Male; **F**: Female), and age (in years) followed by details of the individual implantation scheme, including for each of the 19 patients analyzed for this study, the total number of implanted multielectrodes and the number of contacts placed on each specific implanted brain region.

| Patient Code | Gender (M/F) | Age (years) | Number of Multielectrodes | Number of Contacts | Implanted Regions (Number of contacts) |  |
| --- | --- | --- | --- | --- | --- | --- |
| 1785 | M | 25 | 10 | 71 | Right Calcarine sulcus (1)<br>Right Cuneus (1)<br>Right Lingual gyrus (6)<br>Right Superior Occipital gyrus (3)<br>Right Middle Occipital gyrus (3)<br>Right Inferior Occipital gyrus (10) | Right Fusiform gyrus (5)<br>Right Angular gyrus (6)<br>Right Middle Temporal gyrus (13)<br>Right Inferior Temporal gyrus (8)<br>Right crus I of cerebellum (3)<br>No encephalic regions (12) |
| 1817 | M | 19 | 12 | 69 | Right Middle Frontal Gyrus, Orbital Part (1)<br>Left Hypocampus (2)<br>Left Calcarine sulcus(1)<br>Left Cuneus (1)<br>Left Lingual gyrus (2)<br>Left Middle Occipital gyrus (7)<br>Left Inferior Occipital gyrus (1) | Left Fusiform gyrus (8)<br>Left Angular gyrus (2)<br>Left Precuneus (1)<br>Left Superior Temporal gyrus (3)<br>Left Middle Temporal gyrus (8)<br>Right Middle Temporal gyrus (1)<br>Left Inferior Temporal gyrus (7)<br>Non-encephalic regions (24) |
| 1998 | F | 31 | 12 | 82 | Left Precentral gyrus (5)<br>Right Middle Frontal gyrus, Orbital Part (1)<br>Left Inferior Frontal gyrus, pars opercularis (3)<br>Right Precentral gyrus (7)<br>Left Superior Frontal gyrus (2)<br>Right Superior Frontal gyrus (3) | Left Superior Frontal gyrus, orbital part (8)<br>Right Superior Frontal gyrus, orbital part (9)<br>Left Middle Frontal gyrus (13)<br>Right Middle Frontal Gyrus (10)<br>Left Middle Frontal Gyrus, orbital part (2)<br>Non-encephalic regions (19) |
| 2006 | F | 32 | 7 | 50 | Left Inferior Parietal Gyrus (5) | Left Angular Gyrus (1) |

|  |  |  |  |  | Right Inferior Parietal Gyrus (7) | Right Angular Gyrus (7) |
| --- | --- | --- | --- | --- | --- | --- |
| Code | Gender (M/F) | Age (years) | Number of Multielectrodes | Number of Contacts | Implanted Regions (Number of contacts) |  |
| 2033 | M | 28 | 12 | 79 | Left Insula (1)<br>Left Parahippocampal gyrus (22)<br>Left Fusiform Gyrus (6)<br>Left Superior temporal gyrus (20) | Left Middle Temporal gyrus (21)<br>Left Inferior Temporal gyrus (2)<br>Non-encephalic regions (7) |
| 2040 | M | 29 | 9 | 54 | çLeft Precentral Gyrus (4)<br>Left Supplementary Motor area (8)<br>Left Medial Frontal Gyrus (5)<br>Left Superior Frontal Gyrus (12)<br>Left Anterior Cingulate gyrus (2)<br>Left Middle cingulate gyrus (4) | Left Paracentral Lobule (4)<br>Left Middle Frontal Gyrus (10)<br>Non-encephalic regions (5) |
| 2067 | M | 19 | 11 | 82 | Right Inferior frontal gyrus, pars opercularis (2)<br>Right Inferior frontal gyrus, pars triangularis (2)<br>Right Insula (2)<br>Right Hippocampus (3)<br>Right Parahippocampal gyrus (3)<br>Right Middle Occipital gyrus (1)<br>Right Middle temporal Pole (2) | Right Inferior Occipital gyrus (2)<br>Right Fusiform Gyrus (6)<br>Right Supramarginal gyrus (5)<br>Right Angular gyrus (2)<br>Right Superior Temporal Gyrus (1)<br>Right Middle Temporal gyrus (7)<br>Right Inferior Temporal gyrus (21)<br>Non encephalic regions (23) |
| 2074 | F | 20 | 11 | 66 | Right Superior frontal gyrus, orbital part (1)<br>Left Middle Frontal Gyrus (2)<br>Right Middle Frontal Gyrus (3)<br>Right middle frontal gyrus, orbital part (1)<br>Right Inferior Frontal gyrus ,pars triangularis(1)<br>Left Inferior frontal gyrus, orbital part (1) | Left Olfactory cortex (3)<br>Left Insula (3)<br>Right Insula (13)<br>Right Parahippocampal gyrus (3)<br>Non encephalic regions (11) |
| 2090 | F | 22 | 7 | 48 | Right Superior Frontal gyrus (14)<br>Right Middle Frontal gyrus (13) | Right midcingulate area (1)<br>Right Inferior Parietal gyrus (2) |

|  |  |  |  |  | Right Supplementary motor area (5)<br>Right Superior Frontal gyrus, medial part (1) | Right Middle Temporal Gyrus (1)<br>Non-ncephalic regions (11) |
| --- | --- | --- | --- | --- | --- | --- |
| Patient Code | Gender (M/F) | Age (years) | Number of Multielectrodes | Number of Contacts | Implanted Regions (Number of contacts) |  |
| 2135 | F | 25 | 9 | 75 | Right Precentral gyrus (2)<br>Right Superior Frontal Gyrus, orbital part (1)<br>Right Middle Frontal gyrus (1)<br>Right Middle Frontal gyrus, orbital part (2)<br>Right Inferior Frontal gyrus, orbital part (2)<br>Left Hippocampus (4) | Left Fusiform Gyrus (4)<br>Left Middle Temporal Gyrus (10)<br>Left Middle Temporal Pole (9)<br>Left Inferior Temporal (17)<br>Left lobule III of cerebellar hemisphere (1)<br>Non-encephalic regions (15) |
| 2171 | F | 22 | 8 | 56 | Right Superior Frontal Gyrus (13)<br>Right Middle Frontal Gyrus (17)<br>Right anterior Cingulate area (5) | Right Middle Cingulate area (9)<br>Right Parahippocampal gyrus (4)<br>Non-encephalic regions (8) |
| 2206 | M | 27 | 8 | 50 | Left superior frontal gyrus (6)<br>Left Middle Frontal gyrus (10)<br>Left Middle Cingulate area (2)<br>Left Parahippocampal gyrus (4)<br>Left Postcentral gyrus (3) | Left superior Parietal gyrus (5)<br>Left Precuneus (7)<br>Left Paracentral Lobule (4)<br>Non-encephalic regions (10) |
| 2222 | F | 25 | 8 | 63 | Left Parahippocampal gyrus (27)<br>Left Fusiform gyrus (3)<br>Left Superior temporal gyrus (11) | Left Middle Temporal gyrus (5)<br>Left Inferior Temporal gyrus (6)<br>Non-encephalic regions (11) |
| 2230 | M | 19 | 8 | 60 | Right Precentral gyrus (1)<br>Right Insula (3)<br>Right Parahippocampal gyrus (14)<br>Right Inferior Parietal gyrus (2) | Right Supramarginal gyrus (7)<br>Right Superior Temporal gyrus (19)<br>Right Middle Temporal gyrus (6)<br>Non-encephalic regions (8) |
| 2256 | F | 18 | 10 | 54 | Right Precentral gyrus (7) | Right Middle Cingulate area (3) |

|  |  |  |  |  | Right Superior Frontal gyrus (11)<br>Right Middle Frontal gyrus (14)<br>Right inferior frontal gyrus, pars triangularis (9) | Right Parahippocampal gyrus (3)<br>Right Paracentral Lobule (1)<br>Non-encephalic regions (3) |
| --- | --- | --- | --- | --- | --- | --- |
| Patient Code | Gender (M/F) | Age (years) | Number of Multielectrodes | Number of Contacts | Implanted Regions (Number of contacts) |  |
| 2259 | M | 20 | 8 | 62 | Left Superior Frontal gyrus (1)<br>Left Middle Frontal gyrus (2)<br>Left inferior frontal gyrus,pars triangularis (7)<br>Left Insula (2)<br>Left Anterior Cingulare area (2) | Left Parahippocampal gyrus (30)<br>Left Caudate nucleus (2)<br>Left Superior Temporal gyrus (3)<br>Left Middle Temporal gyrus (4)<br>No- encephalic regions (9) |
| 2270 | F | 22 | 7 | 62 | Left Parahippocampal gyrus (23)<br>Left Superior Temporal gyrus (6) | Left Middle Temporal gyrus (18)<br>Left Inferior Temporal gyrus (4)<br>Non-encephalic regions (11) |
| 2306 | F | 25 | 6 | 48 | Left Midle Frontal gyrus ( 3)<br>Left Parahippocampal gyrus (39)<br>Left Superior Temporal gyrus (3) | Left Middle Temporal gyrus (15)<br>Left inferior Temporal gyrus (13)<br>Non-encephalic regions (8) |
| 2316 | F | 27 | 10 | 74 | Right Insula (1)<br>Left Parahippocampal gyrus (5)<br>Right Parahippocampal gyrus (40)<br>Right fusiform (6)<br>Right Superior Temporal (5) | Left Middle Temporal (2)<br>Right Middle Temporal (18)<br>Right Inferior Temporal (13)<br>Non-encephalic regions (4) |

Each patient was implanted with an average of 9±2 depth electrodes (Adtech, Racine, Wisconsin, USA) and 63±12 contacts (∼7 contacts per electrode). Implantation sites were selected exclusively on clinical criteria, unrelated to the aims of analyses planned for the present study. patients were medication-free at the time intracranial stimulation and iEEG recordings were performed. The implantation procedure was guided with a Leksell stereotactic frame (Elekta, Stackholhom, Sweden) using T1 magnetic resonance imaging sequences (3T, General Electrics, Fairfield, Conneticut, USA) performed prior to the implantation procedure (Figure 1). For each patient, a post-implantation CT scan was co-registered with the normalized T1 MRI sequence in MNI space. The MNI coordinates of each contact were recovered automatically using in-house custom-made software (EPILOC toolbox) developed by the STIM facility (Stereotaxy: Techniques, Images, Models) operating on the CENIR platform at the *Institut du Cerveau et la Moelle Epinière* (Paris, France).

The coordinates of each multielectrode contact were associated to a brain region of the Automated Anatomical Labeling (AAL) human brain atlas^20^. This identification was performed by overlaying MNI normalized contact coordinates to the AAL atlas template using the MRIcron software ^21^. Our analyses included iEEG datasets from electrodes implanted in 66 different anatomical areas, vastly covering left and right frontal, temporal, parietal, occipital regions, as well as the hippocampus (Figure 2). Signals from contacts implanted in the cerebellum, the ventricular system or in structures such as, the skull, meningeal spaces and coverings were not included in our analyses.

**Figure 2.**
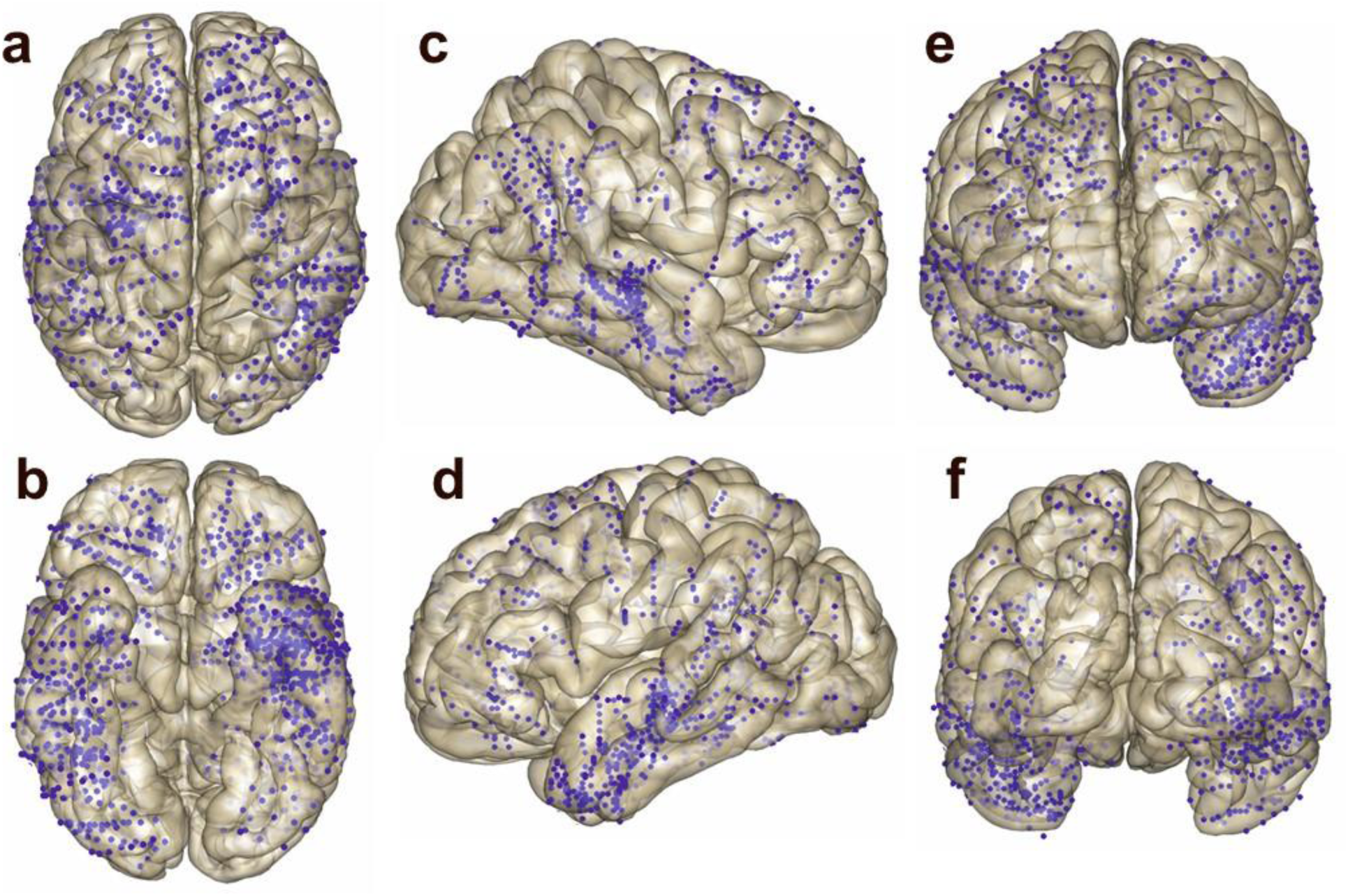
Global multielectrode implantation scheme. Detailed distribution of multielectrode contacts for all patients (n=18) displayed in common normalized MRI brain volume space (MNI 152 coordinates) in Top, Bottom, Left, Right, Rostral and Caudal views. Blue dots represent the projection of each contact site onto the cortical surface of the brain

### Intracranial stimulation and intracranial EEG acquisition procedures

Raw EEG data were recorded continuously from each implanted contact using a 16 bits Micromed Amplifier system (Micromed, Mogliano Veneto, Italy). The sampling rate was set at 1024 Hz and signals were bandpass filtered at 0.15-350 Hz. An external electrode located on the scalp FCz position (10/20 system) was used as a reference for iEEG recordings. During this session, electrical stimulation was performed as a part of the routine clinical protocol put in place to identify the localization of epileptogenic areas ^22^ with a programmable clinical Micromed stimulator. Patients laid comfortably on a bed and were asked to keep their gaze on a fixation cross displayed on the center of a computer screen placed in front of them during the delivery of each stimulation burst. For every patient, each pair of adjacent contacts (within the same multielectrode) was used to systematically deliver bursts made of five singly triggered biphasic pulses (1 ms width) discharged every second.

Pulse shape, pulse duration and pulse polarity (positive wave first) were kept constant across all stimulation sites and studied patients. Nonetheless, for ethical reasons, the stimulation protocol remained exclusively driven by clinical criteria independently of the scientific goals of the present study and could not be subjected to any modification. A customized stimulation approach was applied to each patient guided by the criteria of an expert epileptologist. This procedure aims to causally probe using the least number of electrical pulses, potentially relevant epileptogenic foci and resolve if locations are prone or not to develop epileptiform activity in response to stimulation. During stimulation sessions leading to our dataset, pulses were delivered at increasing intensities (from low-to-high intensity) ranging from 0.5 mA to a maximum of 5 mA, totaling an overall number (across patients) of 3475 pulses (225 pulses at 0.5 mA, 815 pulses at 1mA, 1270 pulses at 2mA, 770 pulses at 3mA, 280 pulses at 4mA, 115 pulses at 5mA). Each subject received an average of 200 pulses (a minimum of 40 and a maximum of 340 pulses). The mean duration of a stimulation session was ∼4 h, including all the resting periods that each patient needed during the session.

### Intracranial EEG data pre-processing

Intracranial EEG responses to single-pulse electrical stimulation were recorded by multielectrode contacts of the implantation scheme not involved in delivering electrical current. For each stimulation pulse we extracted 1400 ms of iEEG data (500 ms prior and 900 ms following the onset of each electrical pulse). Data were pre-processed with a pipeline based on in-house programmed software (Matlab, Mathworks, MA, USA) including an artifact removal procedure to eliminate stimulation artifacts^16^ and a Laplacian data transformation to estimate local field potentials induced by each electrical pulse^23^. Importantly, only single pulse iEEG data from contacts corresponding to areas classified by an expert epileptologist as non-epileptogenic (i.e., documented as ‘healthy’ in a clinical report employed later to discuss and plan neurosurgical approaches) were included in our analyses. Additionally, trace-by-trace and subject-by-subject visual inspection with *ad hoc* help from an expert epileptologist allowed further verification that the selected iEEG dataset were clean of interictal or post-pulse epileptogenic activity and free of any additional sort of artifact (e.g., malfunctioning contacts, 50 Hz noise, contacts placed in non-neural tissue such as subdural spaces or air) which could eventually affect analyses and bias findings. Dataset obtained from sites classified as epileptogenic and eventually considered as potential candidates for neurosurgical resection were left aside and remained unexamined.

Electrical currents injected by the stimulation induced a stereotyped artifact characterized by high-amplitude waveforms, which affected recordings for at least 7 milliseconds ^24^. These were eliminated by removing a 12 ms epoch of the signal (spanning from 1 ms prior to pulse onset to 11 ms thereafter) and interpolating blank periods with a weighted cubic spline interpolation method^16, 22, 25^. This same method was applied to iEEG traces recorded from every contact of the implantation scheme (i.e. each multielectrode from every patient) not involved in delivering the electrical pulse. A composite scheme using the nearest electrode contact neighbor (in the same multielectrode) was applied to estimate the local field potential spatial derivative (i.e., the Laplacian) using the following equation for 1-dimensional implants:

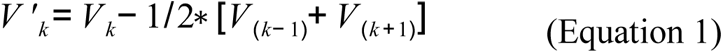

where V_k_ is the recorded field potential at the k^th^ position in the electrode, and V’_k_ is the transformed signal re-referenced to a local reference. Contacts on the upper and deeper portions of the electrode strip were re-referenced to the signal of the closest contact. Using this transformation, we ensured an optimal representation of reliable local signals available from our implantation scheme (see ^23^, for review).

### Time-frequency analyses of iEEG and statistical strategies

Unless indicated otherwise, all analyses were planned *a priori* to substantiate specific stated hypotheses or predictions. The goal of these analyses was to map within a normalized brain volume the local region-specific oscillatory responses induced by single pulses of intracranial stimulation. We sought to identify and characterize the oscillatory signature generated on each stimulated region. This analysis was constrained to AAL atlas regions hosting at least 3 multielectrode contacts: 2 adjacent contacts in charge of delivering the electrical pulse and a 3^rd^ contact able to record local iEEG responses to stimulation (Figure 3A).

**Figure 3.**
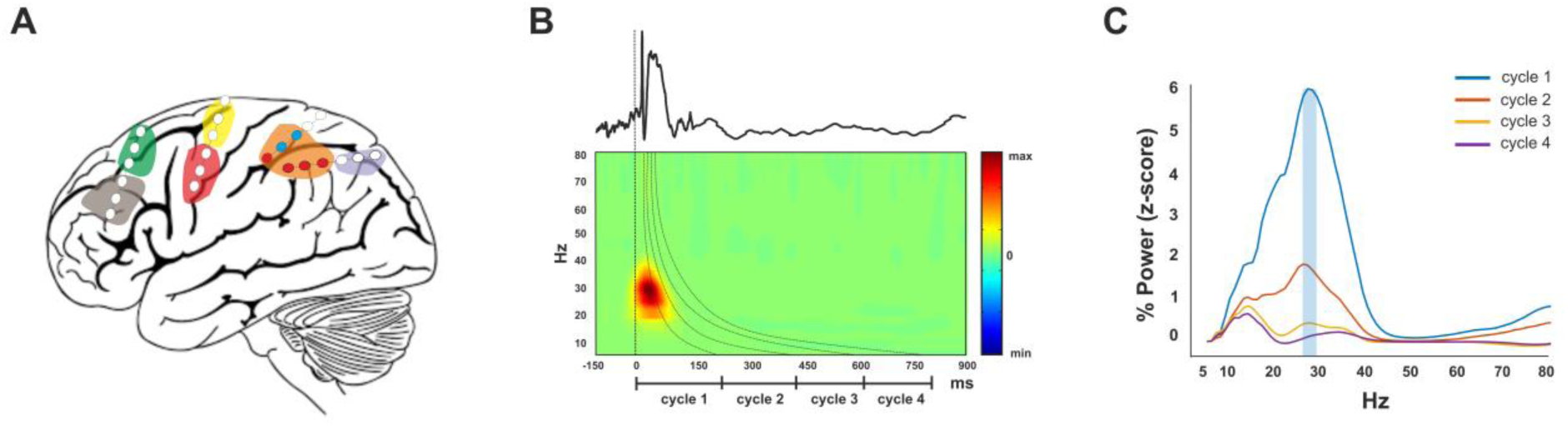
Intracranial EEG analyses of spectral fingerprints and atlasing procedure. (**A**) Schematic drawing showing an example of multielectrode contacts placed in several brain areas (in these case 6 colored regions of the AAL atlas, 4 in the frontal and 2 in the parietal lobe). Blue color signaled contacts delivering the electric pulses whereas red contacts (located in the same region) indicate contacts of the same atlas region used to record iEEG responses to stimulation. In our study we analyzed and averaged through local responses for stimulation and recording contacts located in 33 regions of the AAL atlas. (B, top) Representative example of a iEEG trace corresponding to the mean average response of electrical stimulation pulses delivered to a brain AAL atlas region; (B bottom) Time-frequency analysis of local responses to the single pulse stimulation of the prior iEEG trace. The vertical dashed line signals the time onset of the electrical pulse (t=0). The color-coded scale represents increases (warm hues) and decreases (cold hues) of oscillation power. Dashed lined curved profiles demarcate the length of an oscillation cycle at each frequency bin for the spectral range considered in our analyses (from 5 to 80 Hz, in 1 Hz bins). Note that the baseline pre-stimulation period was adapted to each frequency bin and set to a length of 2 cycles (two periods) oscillation; (C) Representative example of a local spectral fingerprint from a specific cortical site in response to single pulse stimulation. It shows the distribution across frequency bands (x horizontal axis) of power enhancements (y vertical axis) induced by a single electrical pulse during the 1^st^, 2^nd^, 3^rd^ or 4^th^ oscillation cycle (coded in 4 colors) at each frequency bin. The ‘natural’ frequency (indicated with a light blue semitransparent box) was estimated as the frequency bin showing the highest level of power increase with respect to baseline power levels, considering only the 1^st^ oscillation cycle (blue power profile)

Local iEEG responses to intracranial stimulation pulses were analyzed through a three-step process: First, on each patient and for each stimulation event (delivered to the same AAL atlas region), we averaged any available single-trial local iEEG responses recorded by contacts implanted in the same region. Following this procedure, for each patient dataset we obtained a map of region-specific iEEG responses to each stimulation event. Second, we averaged responses across stimulation events delivered to the same AAL atlas region at different stimulation intensities within each patient. This resulted in an accurate region- and patient-response estimate to single pulses of electrical stimulation. Third, we gathered in normalized brain atlas space, local regional- and intensity averaged-iEEG responses to electrical pulses delivered from all patients. As a result, we obtained 33 different datasets (one per each AAL atlas region for which we had stimulation data provided by at least a patient), corresponding to the local average of iEEG responses to each pulse delivered across our cohort of 18 patients.

The averaging of physiological responses to intracranial stimulation between and across individuals is justified by 3 main reasons: First, in order to keep stimulation sessions short, the number of pulses delivered through each pair of contacts into a given site for each individual patients was rather scarce (5 single pulses for each stimulation event (i.e. contact pair, brain site and current intensity), determined by a standardized clinical protocol). Therefore, grouping datasets recorded by electrodes located within the same AAL region and considering each local iEEG response to individual stimulation pulses an independent event proved absolutely necessary to optimize signal-to-noise ratio. Second, electrode implantation schemes were determined by a multidisciplinary team of clinical experts working in the field of epilepsy (neurologists, clinical neurophysiologists, neuroanatomist, and neurosurgeons), guided by clinical requirements considering the unique needs of each patient. Clustering datasets from contacts across different patients on the basis of AAL atlas parcellations allowed us to associate responses from different individuals to specific brain regions. Third, for ethical reasons, the selection of stimulation intensities used on a given patient, on a given site or in a given trial relied exclusively on clinical and neurophysiological criteria applied on a case-by-case basis by an expert epileptologist in charge of stimulation sessions, ignoring any non-clinical consideration.

Data obtained at different intensity levels and considering an unequal number of repetitions (at each stimulation intensity) were used from different brain sites of the same patient across distinct patients. A recent study by our group in this same clinical model, showed that regardless of stimulation intensity, intracranial stimulation (with 50 Hz bursts) induced non-statistically significant increases of gamma power ^16^. We hence assumed that stimulation intensity would not significantly impact the magnitude of power enhancements generated by electrical pulsed perturbations. Accordingly, a representative local estimate of stimulation responses (Figure 3B, top) was calculated by averaging, within patients, different stimulation intensities delivered through a fixed each pair of contacts on the implanted brain location.

To analyze iEEG responses to stimulation in the frequency domain, we convoluted EEG datasets obtained from each AAL atlas region to a complex Morlet wavelet (see ^16^ for details on the procedure and its validation). The analyzed frequencies ranged from 5 to 80 Hz, with a linear increase in 1 Hz in steps. Since due to the need to remove electrical pulse artifacts we interpolated 12 ms intervals of post-stimulus data, we had to exclude unreliable analyses of frequencies higher than 80 Hz. The time-varying energy maps, measured as the square of the absolute value of the Morlet coefficient (a.k.a. power), were computed for each trace from each region-specific iEEG dataset. We then calculated for each 1 Hz frequency bin of the frequential spectrum the percent change in power with respect to a baseline epoch, prior to the onset of the electrical pulse. The baseline period considered for spectral analyses was adapted to each frequency, hence set to a duration of 2 cycles (2 periods of oscillation) for each 1 Hz frequency bin.

Moreover, since iEEG responses were recorded from different subjects and obtained under different stimulation intensities, time-varying power increases at each frequency bin were transformed to z-scores, obtaining a single time-frequency chart characterizing power increases in response to single electrical pulses for each AAL sampled region (See Figure 3B). These normalized time-varying maps were averaged across pulses, to obtain the mean time-varying map of power increases of each region.

Spectral fingerprints of local iEEG responses to single electrical pulses (delivered to each cerebral region) were extracted by measuring the distribution of power enhancements during a time period comprising the 1st cycle of oscillation at each frequency bin (across a selected frequency spectrum spanning from 5 to 80 Hz). For each regional spectral fingerprint, we determined the local *‘natural’ frequency*, which we defined as the frequency bin showing the greatest enhancement of power (i.e., the largest z-value) across the whole fingerprint (Figure 3C).

We also characterized *spectral fingerprint complexity* according to the number of local increases (i.e., peaks or modes) of power between 5 Hz and 80 Hz. In order to consider an increase of power at a given frequency peak, as being reliable and worth trusty, two constraints were imposed: (1) The identified frequency peak had to account for at least 20% of the maximum power of the ‘natural’ frequency band; (2) Two different frequency peaks had to be separated by at least 8 x 1 Hz-frequency bins.

We tested across regions the relationship between the complexity of each spectral fingerprint following stimulation (measured as the number of modes present in their frequency distribution profiles) and the ‘natural’ frequency determined for each site. To this end, ‘natural’ frequencies were categorized in four different bands: *Low Beta* (12-19 Hz)*, High beta* (20-29 Hz)*, Low gamma* (30-39 Hz) and *High gamma* (>40 Hz). Since our results showed that all regional spectral fingerprints elicited after electrical stimulation exhibited ‘natural’ frequencies higher than 12 Hz we did not consider alpha [8-12 Hz] neither theta [4-7 Hz] frequency bands in this analysis. A Chi-squared test was used to determine whether or not the level of complexity of a spectral fingerprint was influenced by the frequency level of the ‘natural’ frequency on each explored region.

We also hypothesized a specific relationship between the so-called ‘natural’ frequency (defined as the frequency showing the maximum increase of power, likely by phase alignment of the local oscillators tied to stimulation) and the cerebral lobe on which stimulation was delivered (see study by Rosanova et al. for further details ^13^). To verify such prediction, we conducted a one-way analysis of variance (ANOVA) with the factor ‘*Cerebral lobe’* (Frontal, Parietal, Occipital and Temporal) as group variable. Additionally, we tested the lobar-specificity of ‘natural’ frequencies induced by intracranial stimulation previously reported by Rosanova et al ^13^, across the above-defined frequency bands as *Low Beta* (12-19 Hz), *High beta* (20-29 Hz), *Low gamma* (30-39 Hz) and *High gamma* (>40 Hz). To this end, a Chi-squared test was used to determine whether or not the frequency band of the ‘natural’ frequency of a given AAL region was associated to each of the 4 cerebral lobes considered in the Rosanova et al. study ^13^.

We also analyzed the duration of the power increases induced by single pulses of intracranial stimulation to each AAL atlas region (specifically at the ‘natural’ frequency). To this end, we averaged power increases at the ‘natural’ frequency for each brain region in 5 different time intervals (adapted to each frequency bin) prior or following the delivery of each electrical pulse (See Figure 3B): *Baseline* (2 oscillation periods at the ‘natural’ frequency) immediately prior to the delivery of the electrical pulse and *1^st^ cycle, 2^nd^ cycle, 3^rd^ cycle, and 4^th^ cycle* (i.e., duration in time of the 1^st^, 2^nd^, 3^rd^ and 4^th^ period of an oscillation at the ‘natural’ frequency) following the delivery of each single electrical pulse. Importantly, these time intervals were tailored according to the oscillation period of the ‘natural’ frequency identified for each anatomical AAL atlas region.

These data were entered into a one-way repeated measures ANOVA with factors ‘*Time interval’* (Baseline, Cycle 1, Cycle 2, Cycle 3, Cycle 4) and *‘Cerebral lobe’* (Frontal, Parietal, Occipital and Temporal). Additionally, we tested if the time course of the power increases at the ‘natural’ frequency differed across frequency bands. To this end, we entered these data into a one-way repeated measures ANOVA with factors ‘*Time interval*’ (Baseline, Cycle 1, Cycle 2, Cycle 3, Cycle 4) and *‘Frequency band’* (Low beta, High beta, Low gamma, High gamma).

Finally, we tested whether the enhancements of power at the ‘natural’ frequency of each region after electrical stimulation were linked to the phase of the ongoing oscillation at these same frequency, at the time single electrical pulses were delivered onto that brain region. With this analysis, we aimed to find a relationship between the level of depolarization vs. hyperpolarization of local neural populations at the time the electrical pulse was delivered (referred to as the ‘up-state’ vs. the ‘down-state’ of an oscillation), and increases of power at the ‘natural’ frequency driven by each stimulation event ^26^.

To this end, for each individual single pulse delivered to a sampled region, we measured the difference between the phase of the ongoing oscillation at the local ‘natural’ frequency immediately before pulse onset (a 12^th^ of cycle before the onset) and the ‘up-state’ of the oscillation (π/2 radians). A non-parametric Spearman correlation test was employed to correlate this relative phase with absolute (non z-scored) levels of power increase at the ‘natural’ frequency during the 1st cycle of the induced oscillation, driven immediately after the delivery of each electrical pulse. The statistical significance of this correlation was confirmed by bootstrapping 10000 samples with replacement from the original data set and estimating the amount of sample values that showed the same sign as the correlation (significance value set as 95% of samples).

Post-hoc tests were conducted with two-tailed t-test between conditions. Degrees of freedom for the ANOVAs were corrected for sphericity (Greenhouse-Geiser ε) when this could not be assumed (Mauchly test of sphericity). For all statistical analyses, significance was set to *p*-values <0.05 and Bonferroni correction for multiple comparisons was applied when necessary.

## RESULTS

### Spatial distribution of ‘natural’ frequencies across brain regions and fingerprint complexity

We successfully determined the spectral fingerprints of power change in response to single pulses of intracranial stimulation for a total of 33 regions of the AAL atlas (Figure 4 & 5). From these we estimated the ‘natural’ frequency, defined as the frequency level showing the greatest enhancement of power following stimulation (Figure 4). On that basis, we then compiled a finer-grained anatomical map of ‘natural’ frequencies gathering and representing data (peak average power of the most predominant frequency from each spectral fingerprint on each sampled AAL region) in a normalized MRI space (Figures 5 & 6).

**Figure 4.**
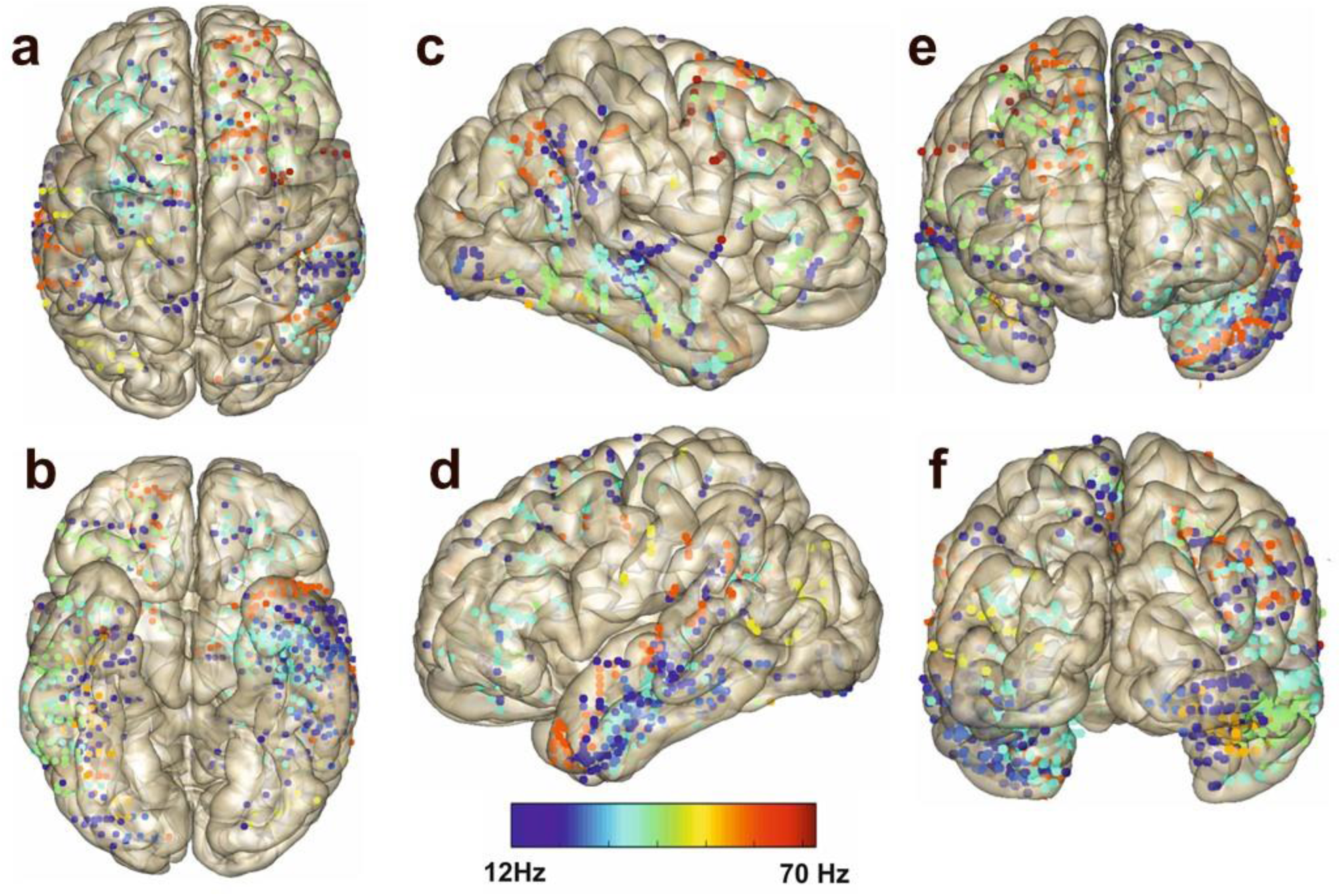
Anatomical distribution of ‘natural’ frequencies in our cohort of patients. Top, bottom, left, right, rostral and caudal views of a normalized MRI brain volume (MNI 152 coordinates) showing in a color-coded scale the location and frequency level (color-coded) of the ‘natural’ frequency at each individual sampled brain site in our cohort of patients. Estimates of site ‘natural’ frequency correspond to the frequency level providing the highest increase of power during the 1^st^ cycle of ‘natural’ oscillatory activity following single pulse stimulation, compared to a pre-pulse baseline. Contacts from each brain region are color-coded to represent the value of the ‘natural’ frequency (12-70 Hz) at that specific location. Cold hues correspond to low frequencies, whereas warm hues correspond to higher frequencies. Data from every multielectrode contact of each patient (n=18) included in our analyses are plotted individually in this figure, regardless of the AAL atlas area they belong to. Notice the complex finer-grained “mosaic-like” spatial distribution of frequency-specific ‘natural’ oscillatory activity across discrete brain sites revealed by this figure

**Figure 5.**
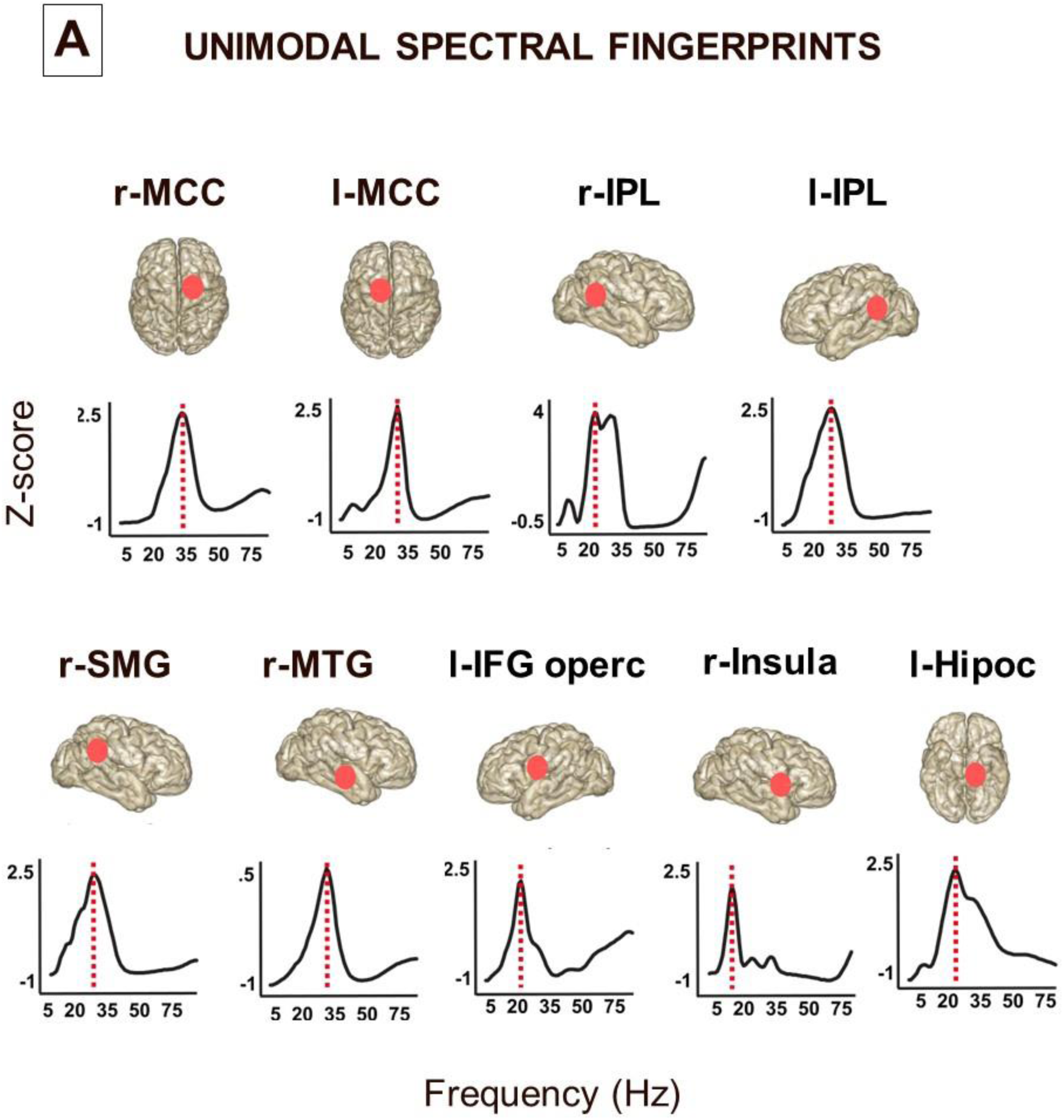
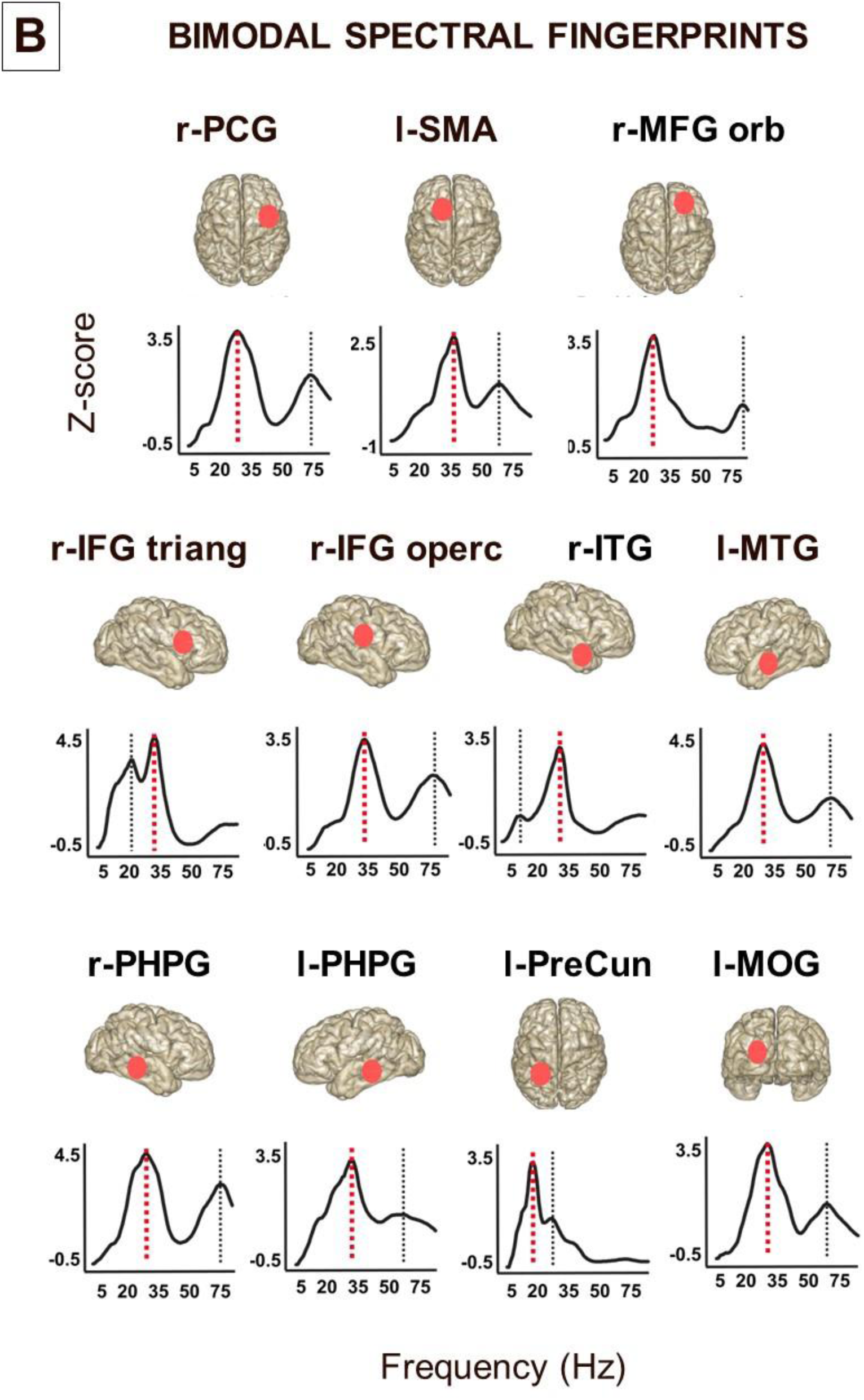
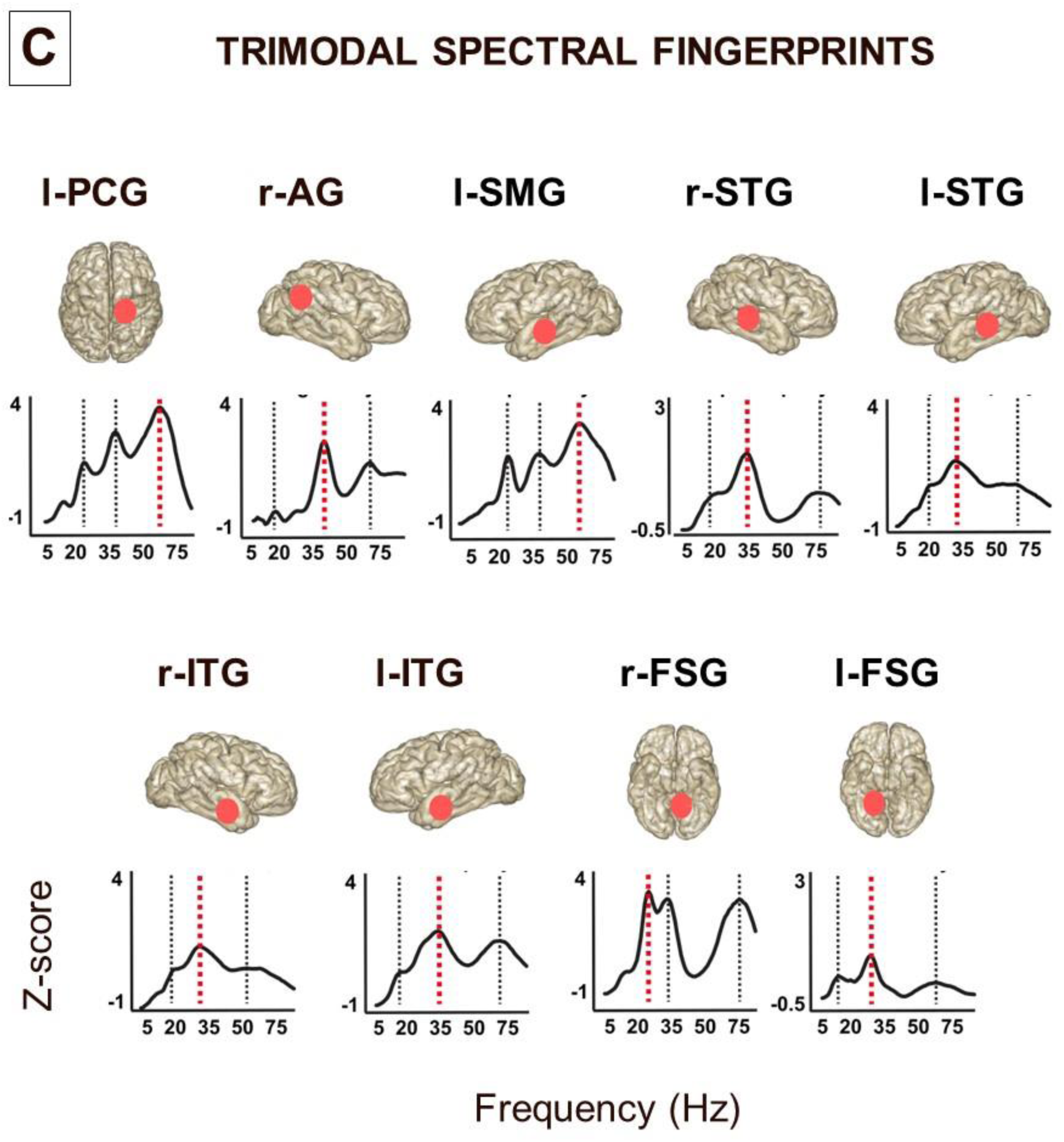
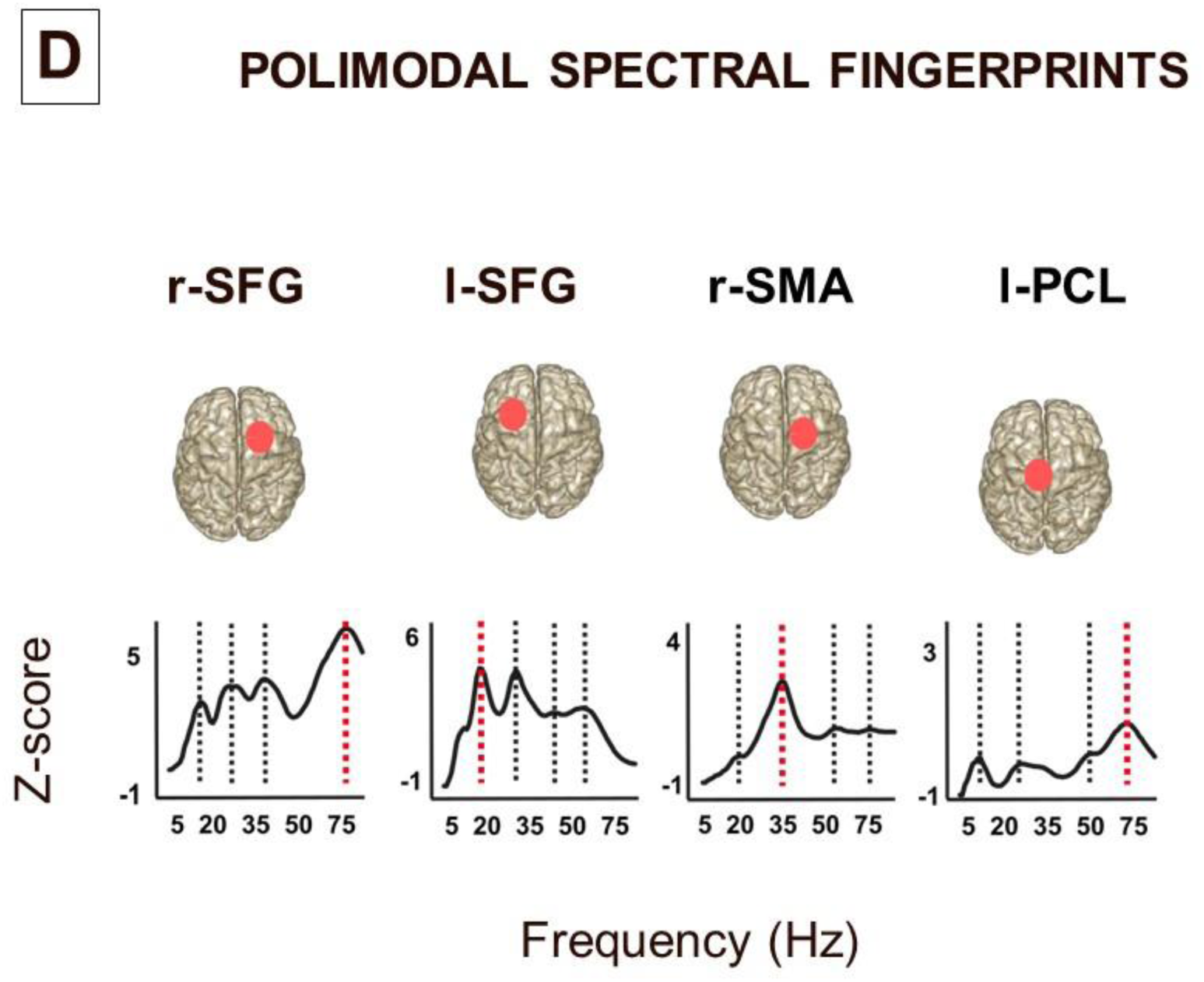
Mean spectral fingerprints of local responses to stimulation for each of the 33 brain regions of the AAL atlas from our cohort of patients. On each individual spectral fingerprint, the red dashed vertical line signals the so called ‘natural’ frequency, i.e., frequency level (between 5 and 80 Hz) showing the maximal peak of power increase, during the 1^st^ oscillation cycle following single pulse stimulation vs. baseline pre-stimulation levels. Black lines signal the presence of additional peaks of power change (from 1 to 4 depending on finger print complexity). Regional fingerprints are presented in 4 separate groups (A, B, C & D) according to their complexity, defined by the number of frequency peaks or modes they contain, characterizing the distribution of power increases across frequency bins of their spectral fingerprints, according to a set of established criteria (see methods for details): Group A: 1 peak; 9 cortical sites, Group B: 2 peaks, 11 cortical sites Group C: 3 peaks, 9 cortical sites and Group D: 4 peaks, 4 cortical sites. In contrast with prior accounts characterizing brain lobes by a single ‘natural’ frequency, note the large diversity and complexity of local spectral fingerprints revealed by our analyses when secondary frequency peaks are considered (Acronyms: **l-MCC**: Left Midcingulate Cortex, **r-MCC**: Right Midcingulate Cortex; **r-IPL**: Right Inferior Parietal Lobule; **l-IPL**: Left Inferior Parietal Lobule; **r-SMG**: right Supramarginal Gyrus; **r-MTG**: Right Medial Temporal Gyrus; **l-IFG**: Left Inferior Frontal Gyrus; **l-IFG-operc**: Left Inferior Frontal Gyrus pars opercularis; **r-IFG-operc**: Right Inferior Frontal Gyrus pars opercularis; **r-Insula**: Right Insula; **l-Hipoc**: Left Hippocampus; **r-PCG**: Right Paracentral Gyrus; **l-SMA**: Left Supplementary Motor Area; **r-MFG-orb**: Right Medial Frontal Gyrus pars orbitalis; **r-IFG** triang: Right Inferior Frontal Gyrus pars triangularis; **r-ITG**: Right Inferior Temporal Gyrus; **l-MTG**: Left Medial Temporal Gyrus; **r-PHPG**: Right Para-Hippocampal Gyrus; **l-PHPG**: Left Para-Hippocampal Gyrus; **l-PreCun**: Left Precuneus; **l-MOG**; Left Medial Occipital Gyrus; **l-PCG**: Left Postcentral Gyrus; **r-AG**: Right Angular Gyrus; **l-SMG**: Left Supramarginal Gyrus; **r-STG**: Right Superior Temporal Gyrus; **l-STG**: Left Superior Temporal Gyrus; **r-ITG**: Right Inferior Temporal Gyrus; **l-ITG**: Left Inferior Temporal Gyrus; **r-FSG**: Right Fusiform Gyrus; **l-FSG**: Left Fusiform Gyrus**; r-SFG**: Right Superior Frontal Gyrus; **l-SFG**: Leftt Superior Frontal Gyrus; **r-SMA**: Right Supplementary Motor Area; **l-PCL**: Left Paracentral Lobule

**Figure 6.**
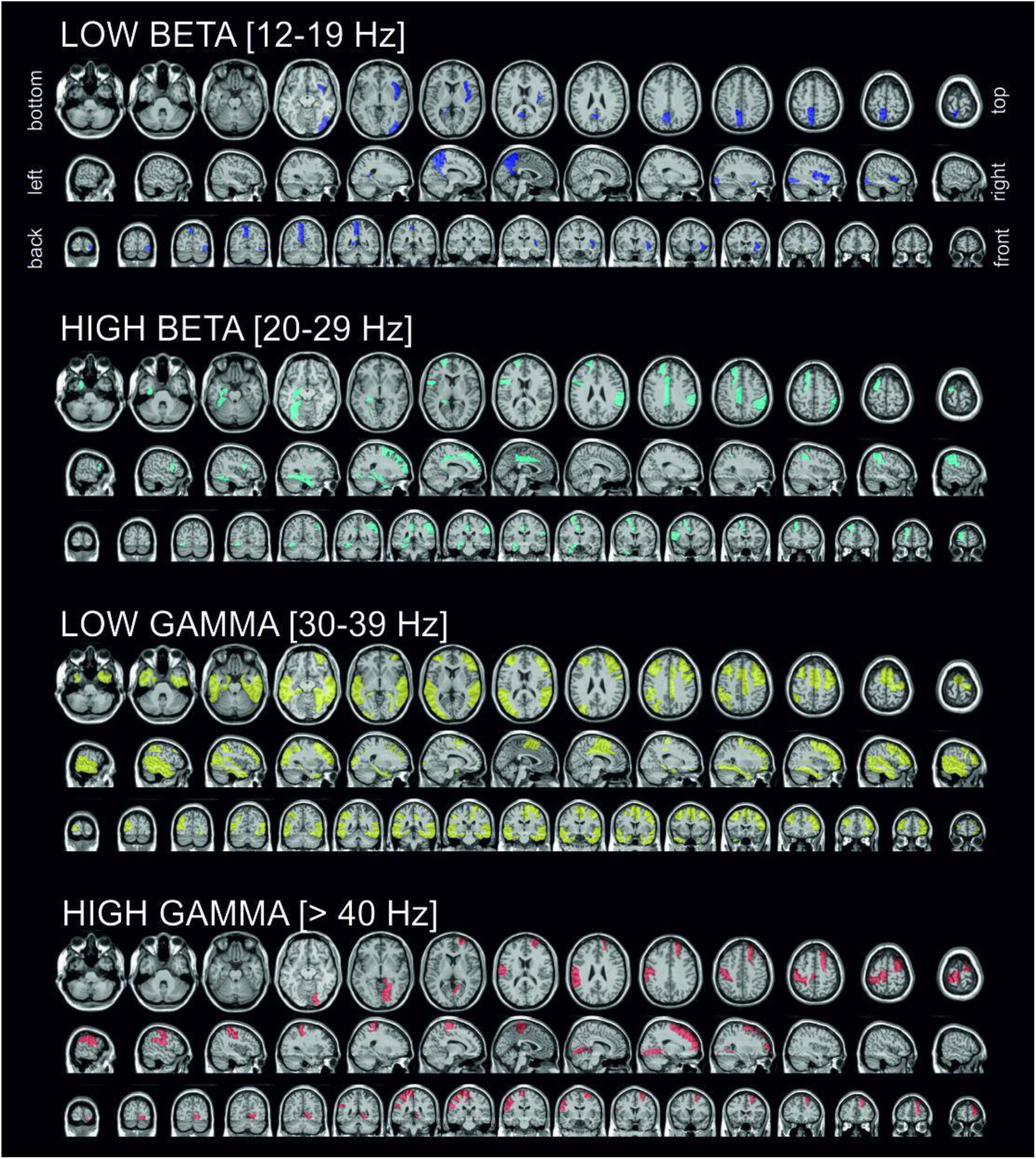
Anatomical distribution of ‘natural’ frequency maps across frequency bands in a human brain atlas. Each local ‘natural’ frequency determined from spectral fingerprints (i.e., highest power change during the 1^st^ cycle post single pulse stimulation vs baseline) were categorized in 4 frequency bands (Low beta: [12-19 Hz]; High beta: [20-29Hz]; Low Gamma: [30-39 Hz]; High Gamma: [>40 Hz]). The final ‘natural’ frequency taken by whole AAL atlas area was defined as the frequency bin displaying the largest enhancement of power during the 1^st^ cycle of the ‘natural’ frequency following single pulse stimulation averaging data from all contacts and patients within each of the 33 AAL areas sampled. Only regions showing their ‘natural’ frequency in one of these 4 frequency band categories (between 12 and 80 Hz) are presented in the figure. Notice ‘natural’ frequencies predominantly at the high-beta and low-gamma bands band in frontal and temporal areas, and within the low gamma band in parietal brain regions.

At difference with prior reports using non-invasive brain stimulation and recording approaches ^13^, no significant relationship was found between local ‘natural’ frequency (frequency showing maximal peak of power across the spectral fingerprint) and cerebral lobe [Lobe effect, F(3,32)=0.59, p=0.623]. However, when ‘natural’ frequencies were categorized in 4 specific frequency bands, a Pearson Chi-square test revealed significant dependency between ‘*Frequency band*’ and ‘*Cerebral lobe*’ [χ^2^(9, N=33) = 17.41, p<0.05], indicating that the proportion of regions showing a ‘natural’ frequency in each of such frequency bands differed significantly across cerebral lobes (see Figure 5 & 6B).

Indeed, in response to single electrical pulses, the majority of AAL regions hosted in the frontal (61.5%) and temporal (83.3%) lobes revealed a maximum power peak at frequencies within the low-gamma band, whereas regions in the parietal lobe reacted differently: 50% of parietal regions showed maximal power increases within the high gamma band and 37.5% of them had responses in the beta band (12.5 % in Low beta band and 25% of regions in High beta band) (Figure 7, Table 2).

**Figure 7.**
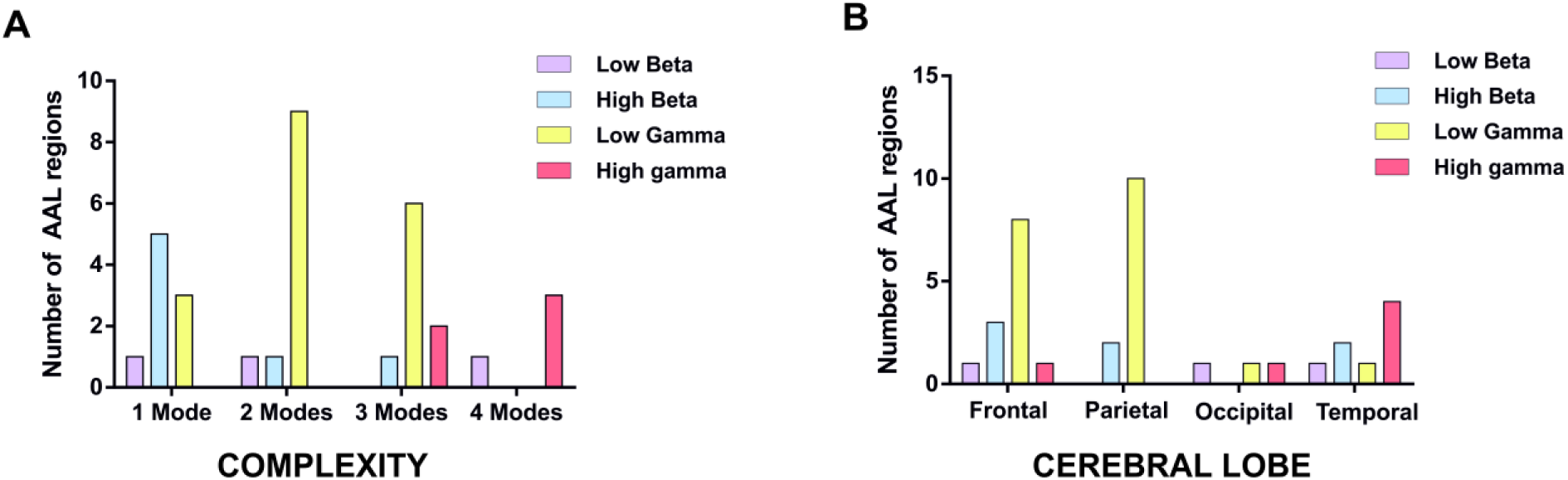
Number of brain regions according to the AAL atlas with a ‘natural’ frequency band. associated to (left panel, A) spectral fingerprint complexity level or (right panel, B) specific brain lobes (frontal, parietal, temporal, occipital). The complexity of the spectral fingerprint was captured by the number of peaks or modes (from 1 to 4) shown by mean regional spectral fingerprint (showing power increases during the 1st oscillation cycle of each frequency following stimulation compared to pre-stimulation baseline). Note that a majority of AAL regions with bi-modal (two peaks) and tri-modal (three peaks) spectral fingerprints (panel A) showed ‘natural’ frequencies within the low-gamma band (80%), whereas the majority of regions showing a unimodal spectral fingerprint (i.e., a single frequency peak) showed ‘natural’ frequencies at the low-beta band (∼60%). Also notice (panel B) ‘natural’ frequencies predominantly in the high-beta and low-gamma bands band in frontal and temporal lobes, and within the low gamma band for AAL regions in the parietal lobe

**Table 2.**
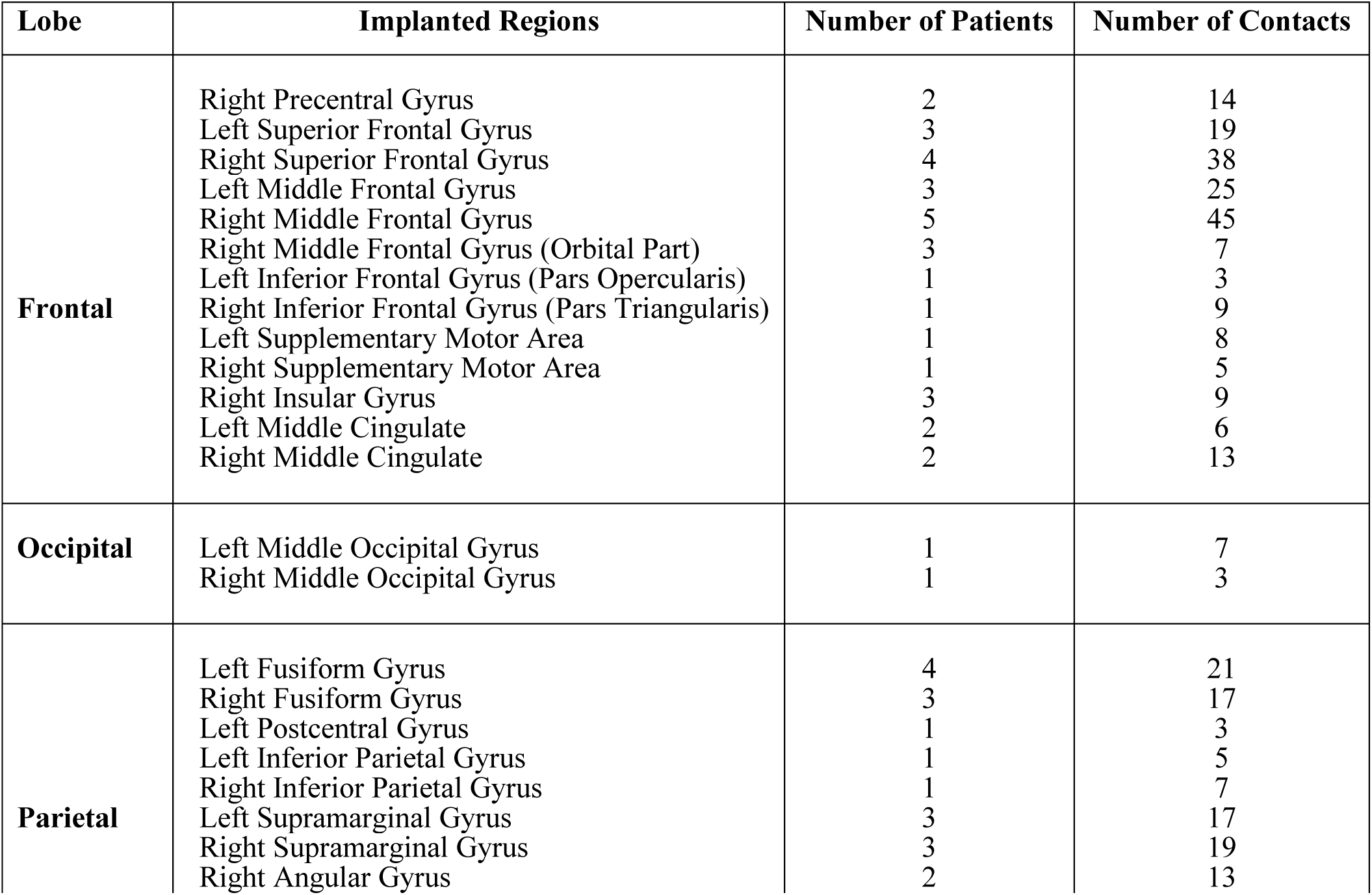
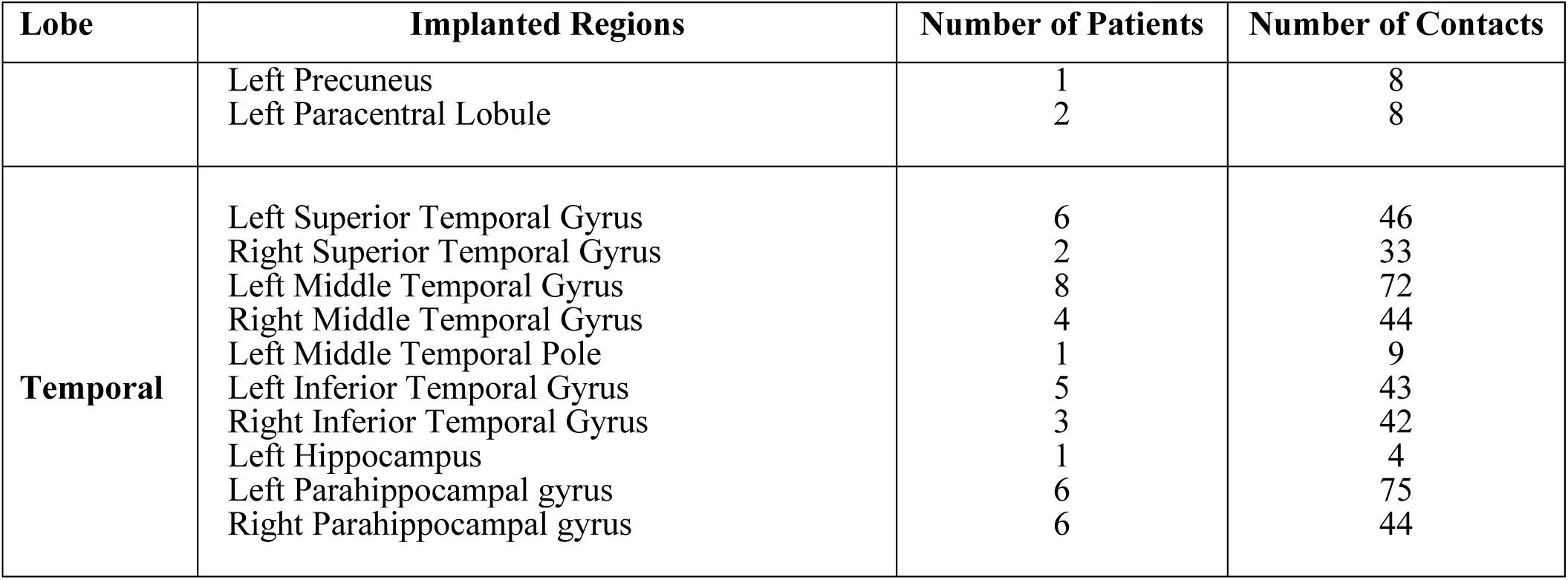
Summary of the implanted regions used to produce the atlas of natural frequencies. These regions are classified by lobe, number of patients and total number of contacts across patients.

| **Lobe** | **Implanted Regions** | **Number of Patients** | **Number of Contacts** |
| --- | --- | --- | --- |
| **Frontal** | Right Precentral Gyrus | 2 | 14 |
|  | Left Superior Frontal Gyrus | 3 | 19 |
|  | Right Superior Frontal Gyrus | 4 | 38 |
|  | Left Middle Frontal Gyrus | 3 | 25 |
|  | Right Middle Frontal Gyrus | 5 | 45 |
|  | Right Middle Frontal Gyrus (Orbital Part) | 3 | 7 |
|  | Left Inferior Frontal Gyrus (Pars Opercularis) | 1 | 3 |
|  | Right Inferior Frontal Gyrus (Pars Triangularis) | 1 | 9 |
|  | Left Supplementary Motor Area | 1 | 8 |
|  | Right Supplementary Motor Area | 1 | 5 |
|  | Right Insular Gyrus | 3 | 9 |
|  | Left Middle Cingulate | 2 | 6 |
|  | Right Middle Cingulate | 2 | 13 |
| **Occipital** | Left Middle Occipital Gyrus | 1 | 7 |
|  | Right Middle Occipital Gyrus | 1 | 3 |
| **Parietal** | Left Fusiform Gyrus | 4 | 21 |
|  | Right Fusiform Gyrus | 3 | 17 |
|  | Left Postcentral Gyrus | 1 | 3 |
|  | Left Inferior Parietal Gyrus | 1 | 5 |
|  | Right Inferior Parietal Gyrus | 1 | 7 |
|  | Left Supramarginal Gyrus | 3 | 17 |
|  | Right Supramarginal Gyrus | 3 | 19 |
|  | Right Angular Gyrus | 2 | 13 |
|  | Left Precuneus | 1 | 8 |
|  | Left Paracentral Lobule | 2 | 8 |
| **Temporal** | Left Superior Temporal Gyrus | 6 | 46 |
|  | Right Superior Temporal Gyrus | 2 | 33 |
|  | Left Middle Temporal Gyrus | 8 | 72 |
|  | Right Middle Temporal Gyrus | 4 | 44 |
|  | Left Middle Temporal Pole | 1 | 9 |
|  | Left Inferior Temporal Gyrus | 5 | 43 |
|  | Right Inferior Temporal Gyrus | 3 | 42 |
|  | Left Hippocampus | 1 | 4 |
|  | Left Parahippocampal gyrus | 6 | 75 |
|  | Right Parahippocampal gyrus | 6 | 44 |

According to the number of local increases (i.e., peaks or modes) of power in bands between 5 and 80 Hz, fingerprints revealed up to four levels of complexity (see Figure 5). Fingerprint complexity was significantly associated with the frequency band of the ‘natural’ frequency on a given region [χ^2^ (9, N=33) = 26.47, p<0.01]. Interestingly, a majority of AAL regions with bi-modal (two peaks) and tri-modal (three peaks) spectral fingerprints had ‘natural’ frequencies within the low-gamma band (80%). At difference, a majority of regions featuring unimodal spectral fingerprints (i.e., a single frequency peak) displayed ‘natural’ frequencies at the low-beta band (∼60%) (Figure 6).

### Duration of power enhancement of induced ‘natural’ frequency across cycles

We here tested whether the duration of power enhancements at the ‘natural’ frequency band on each tested cerebral site lasted beyond the 1st oscillation cycle after each electrical stimulation pulse. We also explored if such duration depended on the frequency band of the ‘natural’ frequency and/or on the cerebral lobe on which they were induced. A main effect of the factor ‘*Time interval*’ [F (4,128) = 79.73, p < 0.001] suggested that power increases at the ‘natural’ frequency triggered by electric stimulation differed across the 1^st^ and consecutive (2^nd^, 3^rd^, and 4^th^) oscillation cycles. Electrical stimulation induced power increases that achieved their maximum during the 1st cycle [t(32) > 9.8, p<0.001 for all the comparisons], and decayed progressively thereafter, extinguishing when oscillations reached the 3^rd^ cycle post electrical pulse [2^nd^ cycle vs. 3^rd^ cycle, t(32) = 10.22, p<0.001; 3^rd^ cycle vs. 4^th^ cycle, t(32) = 2.9, p=0.069] (Figure 8A).

**Figure 8.**
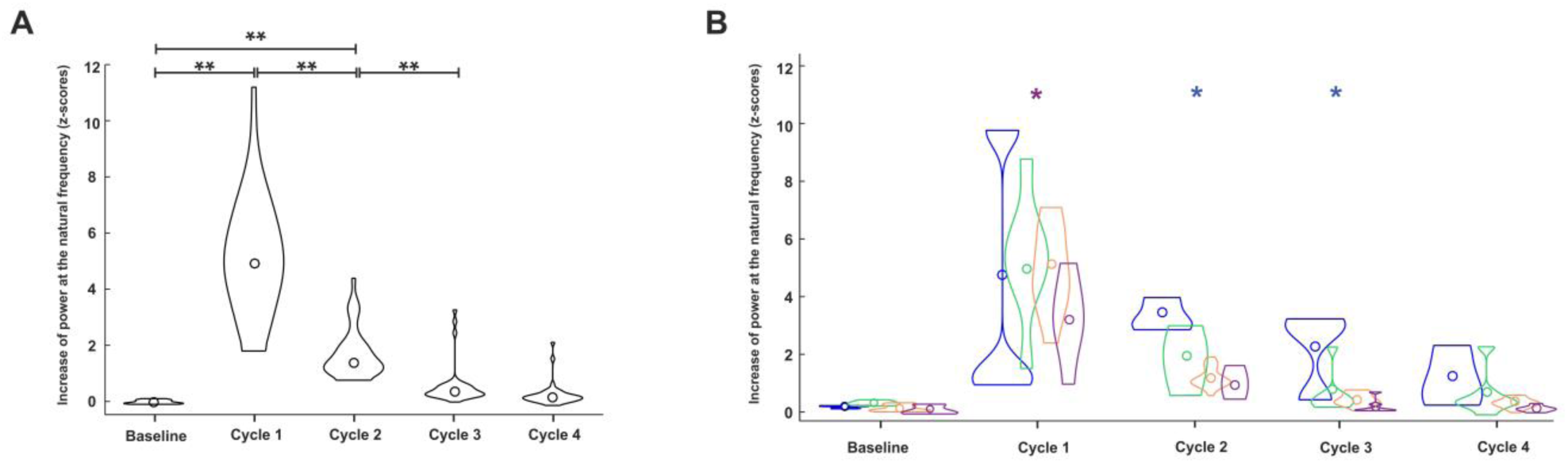
Temporal dynamics of changes in power at the ‘natural’ frequency across four consecutive cycles post-single pulse stimulation. Violin plots corresponding to (left panel, A) Combined average of power changes for all patients (n=19) including all sampled regions and frequency bands and (right panel, B) average of combined power changes from all patients (n=19) including all sampled regions separated by frequency bands (Low beta: [12-19 Hz]; High beta: [20-29Hz]; Low Gamma: [30-39 Hz]; High Gamma: [>40 Hz]). * p < 0.05; ** p < 0.01 vs baseline power levels. Note that the power increases induced by phase alignment are very prominent during the 1^st^ oscillation cycle and decrease quickly as time and cycles elapse (panel A). Brain regions with ‘natural’ spectral responses in the high-gamma band show shorter-lasting power enhancements that the rest, and regions with ‘natural’ frequencies within the low beta band display longer-lasting responses.

No significant interaction between factors ‘*Time interval*’ and ‘*Cerebral lobe*’ [F (4, 128) = 1.73, p = 0.16, ε = 0.33] was found in our dataset, indicating that site-specific temporal dynamics of ‘natural’ frequency power enhancement was not determined by the brain lobe in which these phenomena were being probed. However, factors ‘*Frequency band*’ and *‘Time interval’* [F (4,128) = 1.9, p = 0.045] interacted significantly, supporting an influence of ‘natural’ oscillatory frequency in the temporal dynamics of power modulations across several of their respective consecutive cycles.

Indeed, during the 1^st^ oscillation cycle following stimulation, regions featuring enhanced high-gamma ‘natural’ activity reached significantly lower increases in high-gamma power than regions with ‘natural’ frequencies at any other band [low beta vs. high-gamma, t(7) = 3.56, p=0.03; high beta vs. high gamma, t(11) = 3.1, p= 0.045; low gamma vs. high gamma, t(24) = 2.4, p = 0.025; p>0.05 for the remaining comparisons]. At difference, during the 2^nd^ oscillation cycle following the onset of the electrical pulse, regions with ‘natural’ frequencies in the low beta band showed higher increases of power than the rest [low beta vs. high beta, t(8) = 2.51, p=0.036; low beta vs. low gamma, t(21) = 10.2, p <0.001; low beta vs. high gamma, t(7) = 3.4, p = 0.04; p>0.05 for the remaining comparisons]. Similarly, during the 3^rd^ oscillation cycle, regions with ‘natural’ frequencies in the low-beta band showed a trend towards significantly higher increases of power than those featuring high-beta ‘natural’ activity (low beta vs. high beta, t(8) = 1.99, p=0.062), whereas power increases proved significantly higher for regions featuring low-gamma activity [low beta vs. low gamma, t(21) = 5.9, p <0.001; low beta vs. high gamma, t(7) = 3.16, p = .016; p>0.05 for the remaining comparisons (Figure 8B).

In sum, regions featuring ‘natural’ frequencies in the high-gamma band experienced the lowest increases of power during the 1^st^ oscillation cycle following single electrical pulses, whereas, regions featuring ‘natural’ frequencies in the beta band showed longer-lasting decays of power (higher enhancements of power during the 2^nd^ and 3^rd^ oscillation cycles post pulse) compared to the remaining sampled brain regions.

### Relationship between increase of power at the first cycle and on-going oscillations

Finally, we tested whether absolute increases of power at the ‘natural’ frequency induced by individual electrical pulses were influenced by the phase of the ongoing oscillation at the time of the stimulation pulse. This was accomplished by measuring time differences between the latter and the ‘up-state’ (π/2 radians) phase of the oscillation (Figure 9A). A significant correlation was found between absolute increases of power and the difference between the instantaneous phase and ‘up-states’ at electrical pulse onset (ρ = -0.186; p <0.0001).

**Figure 9.**
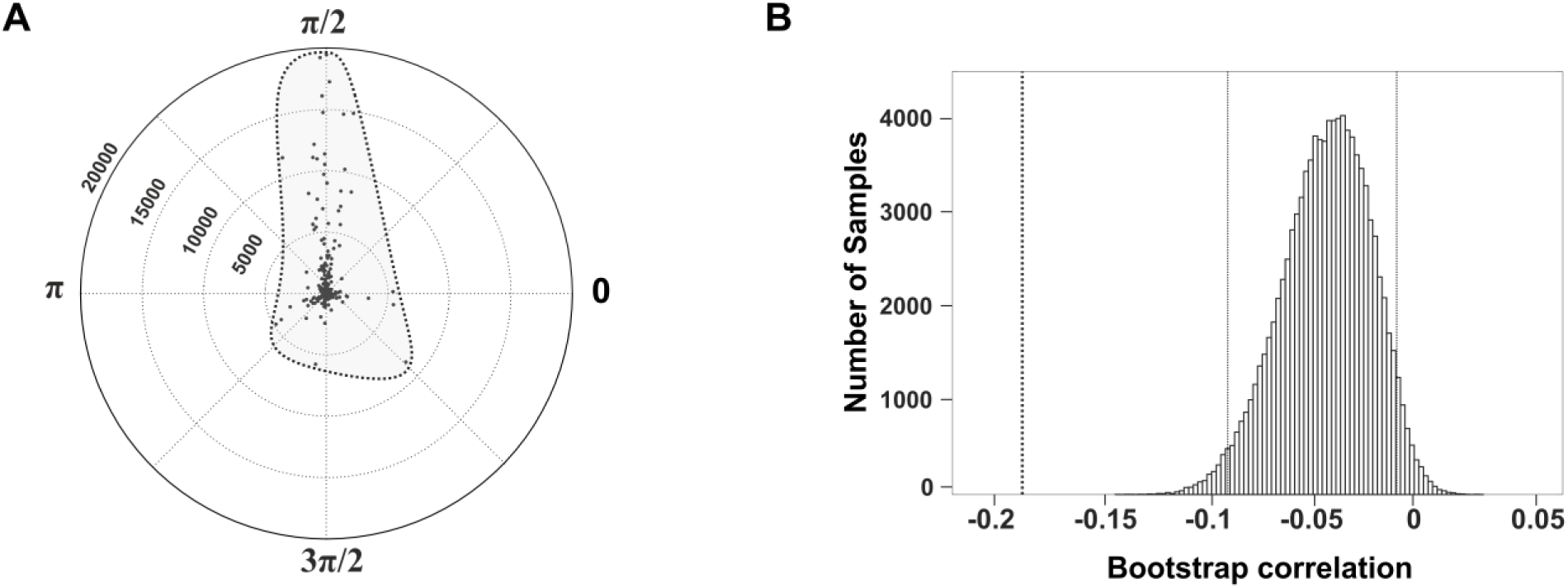
Impact of the ongoing oscillation phase on power increases at the ‘natural’ frequency following single stimulation pulses. (A, left panel) Polar map displaying the magnitude of the power change at the ‘natural’ frequency as a function of the phase of the ongoing oscillation (at this same frequency) at which the pulse was delivered. Notice that pulses delivered in the vicinity of the *‘up-state’* of the ‘natural’ frequency oscillation (π/2 phase) induced higher enhancements of power than those delivered at other phases (0π, π, 3π/2). The black dashed profile signals the linear fit of the correlation; This data supports the phase- and state-dependent nature of natural frequency power enhancements with pulsed electrical perturbations. (B, right panel) Results of the distribution of correlation values after recalculating them by boot-strapping 10000 samples with replacement from the original dataset (1035 single pulses). The bold vertical dashed line indicates the correlation value of the original dataset. The thin vertical black dashed lines signal the 25^th^ and 75^th^ percentile values of the distribution of correlation values obtained with the *boot-strapping* method. Since no correlation value within the boot-strapping distribution exceeds the zero value and less than a 5% of this correlation values were more significant than the correlation value of the original dataset, the correlation is statistically robust and significant.

In order to confirm the statistical significance of this observation, we recalculated the correlation by bootstrapping 100000 samples (with replacement from the original dataset (n=1035)). This approach showed that the sampling distribution of outcomes was skewed negatively (95% confidence intervals between -0.084 and -0.017) (see Figure 9B for details). Since the confidence interval did not include any value greater than 0, we confirmed that the stated correlation was robust and statistically significant.

In sum, increase of power at the ‘natural’ frequency induced by single intracranial pulses delivered on a given region proved strongly dependent from the phase of the ongoing oscillation on that same frequency band.

## DISCUSSION

### Enhancing natural frequencies with single pulse electrical perturbations

Optimal synchronization of ongoing oscillatory activity orchestrated by an external rhythmic input requires both rhythms to cycle with the exact same period ^14^. To fulfill such condition, the external force (for example a rhythmic visual or auditory pattern delivered to the retina or the external ear, or an electrical pulse delivered to a brain area) must adjust the phase of local oscillators, aligning the phases of a diversity of ongoing rhythms operating on the stimulated brain region. Computational models of synchronization predict that among other important parameters (for example waveform amplitude/strength or oscillation duration), the relative frequency of the external stimulation source with regards to the predominant frequency of ongoing oscillations (the so called ‘natural’ frequency) is essential to allow an efficient interaction between extrinsic rhythmic patterns (i.e., imposed by pulsed electrical perturbations) and internal brain rhythms^1^.

In our study, we mapped the intrinsic ‘natural’ frequencies of specific cortical human brain areas using iEEG responses to single electrical pulses. Datasets were obtained from a cohort of implanted epilepsy patients during a clinically-guided mapping quest to identify seizing foci and carried out prior to considering neurosurgical removal of epileptogenic regions. Existing evidence on extrinsically driven modulation of brain oscillations in humans ^10, 14^ supports the notion that brief local perturbations causes the phase-resetting of ongoing EEG rhythms generated by local neural populations, acting as local rhythm generators or oscillators. Because of a progressive alignment in phase, the power of the predominant rhythms generated by such oscillators (the so called ‘natural’ rhythm) is increased. We here hypothesized that such frequency-specific increases of power could serve as proxies of the oscillatory properties operating ‘naturally’ in a locally stimulated brain region.

The results of our study support our ability to identify, map, and characterize the spatial distribution of site-specific spectral fingerprints associated to ‘natural’ oscillatory activity in highly discrete regions of the human brain. Moreover, this evidence could be likely used to infer information about the structural and neurophysiological features of the local circuits (i.e., oscillators) generating such patterns and better understand their contributions to cerebral dynamics and cognitive processing. Equally important, our datasets compiled in an atlas of ‘natural’ spectral fingerprints may provide useful information to optimize the entrainment of local oscillatory activity with invasive (intracranial electrical stimulation) and noninvasive (TMS/tACS) neurostimulation technologies in exploratory, diagnostic or therapeutic applications.

Our analyses showed that singly delivered electric pulses induced site-specific modulations of iEEG activity and revealed the spatial specificity, complexity and diversity of spectral fingerprints in 33 regions of the AAL human brain atlas. Whereas some regions showed a clear increase of power in a specific frequency band (unimodal fingerprints), other brain sites displayed increases in power spanning across contiguous frequency bands with two (bimodal fingerprints) or several power peaks (multimodal fingerprints). The level of spectral fingerprint complexity assessed by the number of modes or peaks generated on a given site depended on the band hosting the ‘natural’ frequency peak and proved higher for high-gamma than for low-beta band activity. Importantly, power increases at the so called local ‘natural’ frequencies started after the delivery of the stimulation pulse and wore off 1.5 to 2 cycles thereafter. Moreover, power changes were tied to the instantaneous phase of ongoing ‘natural’ oscillations fluctuating at the same frequency, at the time single electrical pulses were delivered. This observation supports the hypothesized notion of more efficient enhancement of induced oscillatory power when the onset time of the stimulation pulse is close to the ‘up-state’ of the ongoing ‘natural’ oscillation.

### Mapping natural frequencies in the human brain with intracranial methods

A rich body of literature has reported EEG/MEG evidence in support of power enhancements of local oscillations, immediately following single or multiple perturbations of peripheral sensory stimulation (auditory or visually) or transcranial magnetic/electrical stimulation (TMS/tACS)^10, 21^. Indeed, even if based on a low number of brain regions, past scalp EEG and MEG evidence supports the notion that the physiological response to electrical perturbations ‘echoes’ the spectral features of local ongoing oscillations.

In a seminal study using combined TMS-EEG recordings, magnetic pulses over the left primary motor area modulated generators of alpha and beta oscillatory activity^27^. Similarly, beta band power increases emerging from intrinsic regional circuits have been reported to follow stimulation of the motor cortex^28^. These findings motivated the use of single TMS pulses coupled to high density EEG recordings and served to report consistent enhancements of dominant alpha-band oscillations in the occipital cortex, beta-band oscillations in the parietal cortex and fast beta-gamma band oscillations in the frontal cortex ^13^. The latter outcome suggested that the perturbation of cortical brain regions was followed by common lobar responses at specific dominant frequency bands, mediated by three specific cortico-thalamic modules (involving Brodmann areas 19, 7 and 6, respectively) which reflected the structural and functional organization of stimulated systems.

Our study extends this evidence in three significant directions. First, it provides first-time proof in favor of a complex finer-grained “mosaic-like” spatial distribution of frequency-specific ‘natural’ oscillatory activity induced by singly delivered pulses (Figure 4). Second, it supports the feasibility of combining intracranial stimulation and iEEG recordings from non-epileptogenic regions in cohorts of implanted epilepsy patients to accurately map the distribution of ‘natural’ frequencies across cerebral systems, and regionalize (according to the AAL atlas parcellation) the human brain on the basis of such predominant rhythms. Third and last, our methods improve the spatial resolution achieved by prior pioneering non-invasive stimulation and recording approaches^13^, limiting both, smearing effects and the confounding of volume conduction of scalp EEG recordings.

In a recent study, Frauscher and colleagues ^28^ compiled an atlas of intracranial resting state EEG data obtained in a large sample of epileptic patients, and reported frequency distributions maps across different cortical regions. The mapping outcomes provided here by our analyses based on specific frequency features induced by direct intracranial stimulation reveal complementary features about the distribution of ‘natural’ frequencies throughout cortical brain areas. Yet they also allow comparison with those reported previously with TMS and scalp EEG recording across large lobar regions^13^. To this end, we categorized the frequency spectrum of our study (5 Hz to 80 Hz) in four different bands (Low-beta [12-19 Hz], High-beta [20-29 Hz], Low-gamma [30-39 Hz] and High-gamma [>40 Hz]. Post pulse power modulation effects recorded by each individual contact were grouped across AAL brain atlas regions and were associated to one of the 4 lobes (Frontal, Parietal, Temporal and Occipital Lobe) previously tested for similar purposes using TMS ^13^ (Figure 5 and 6). Our data show that single intracranial electrical pulses delivered to areas within the frontal and temporal lobes induced ‘natural’ frequencies, predominantly in the high-beta and low-gamma bands. This outcome dovetails nicely with the above-mentioned evidence reporting fast beta-gamma band oscillations following frontal TMS stimulation^13^ and resting state intracranial EEG recordings ^28^.

Additionally, resting-state recordings in epileptic patients, during scalp EEG or electro-corticography (ECOG) have also revealed gamma peaks in temporal (entorhinal cortex, parahippocampal gyrus and temporal pole), and frontal (orbitofrontal cortex and the frontal pole) areas^13, 17, 29^. However, the large majority of parietal lobe regions probed in our study with intracranial stimulation revealed ‘natural’ frequencies within the low gamma band, an outcome difficult to reconcile with the beta-band power increases after single pulse TMS reported previously for this same lobe ^13, 29^. Our finding should however not be surprising as gamma rhythms are well-widespread cortically ^30^, can be entrained by stimulation in many brain regions ^16^ and are known to contribute to many cognitive operations and high-level functions. The discrepancy could be simply explained by the frequency cut-off levels considered for analyses (which in Rosanova et al. ^13^ remained below <50 Hz) or by differences in the ability of intracranial vs. transcranial stimulation approaches to phase-reset oscillators in complex “hub regions” such as the posterior parietal lobe. Additionally, Rosanova and colleagues restricted their analysis to a specific region within

Brodmann area 7 in the parietal lobe ^13^, whereas our mapping effort included different regions of the parietal lobe (such as Left Postcentral Gyrus, Left and Right Inferior Parietal Gyrus, Left and Right Supramarginal gyrus, Right Angular Gyrus and Left Precuneus, encompassed by Brodmann area 7, but also 1, 2, 3, 5, 39 and 40). Therefore, sampling biases of the specific subareas of the parietal lobe probed and recorded with methods at very different spatial resolution could also account for such discrepancy.

### Temporal dynamics and phase dependence of natural frequency enhancements

Our study gathered evidence on the dynamics of power enhancements at the ‘natural’ frequency across post-electrical pulse onset time. To this regard, we found maximal increases of oscillation power during the 1st cycle of post-pulse ‘natural’ oscillations, dwindling progressively across the 2nd cycle and wearing off significantly from the 3rd cycle onwards (Figure 8A). The short-lasting nature of single pulse-induced oscillatory enhancement unearthed in our study is in agreement with prior findings reporting entrained rhythmic EEG responses to single or repetitive TMS pulses, as rapidly decaying over time and lasting at most for 1 to 2 oscillation cycles ^8, 10, 28^. We also found that power enhancements across consecutive cycles depended on the frequency band of the ‘natural’ oscillation featured by each given region (Figure 8B). Brain regions with ‘natural’ spectral responses in the high-gamma band showed less pronounced and shorter-lasting power enhancements that the rest. Conversely, regions with ‘natural’ frequencies within the low beta band displayed longer-lasting responses (measured in number of cycles), extending power increases to the 3^rd^ oscillation cycle. Interestingly, this association between frequency and power reported in our study mimics previously reported decreases of power duration, at increasing levels of frequency for electromagnetic brain signals^31, 32^.

In our study, we also report that power increases at the ‘natural’ frequency induced by individual electrical pulses are state-dependent, and accordingly, influenced by the instantaneous phase of the ongoing local oscillation, at the time of the stimulation. More specifically, our analyses showed that pulses delivered during the ‘up-state’ phase (π/2) of an ongoing ‘natural’ oscillation were followed by higher increases in power at this same frequency band than those delivered at any other phase (Figure 9A). As stated elsewhere^13, 14^, this result supports a perturbation-driven phase-resetting mechanistic hypothesis, according to which single stimulation pulses act as an external ‘go signal’ promoting the phase alignment of ongoing rhythms specifically at the natural frequency of the region, and suggest that such phenomenon requires a significant level of ongoing activity prior to pulse onset to occur ^33, 34^.

### Local spectral fingerprint features: diversity and level of complexity

A final crucial aspect that deserves attention concerns the complexity and diversity of the local spectral fingerprints revealed by intracranial stimulation across cortical regions. Indeed, following a focal single pulse electrical perturbation, some regions (parcellated according to the AAL atlas) showed during the 1^st^ oscillation cycle a single unimodal spectral distribution of power enhancement, similar to those previously characterized non-invasively^13^. Nonetheless other cortical areas revealed spectral signatures of higher complexity, made of several power peaks, often spanning across different frequency bands (Figure 5). The constrained scope of our study, based on clinically-guided mapping datasets from implanted patients limits at this time a more profound interpretation of site-specific structural and physiological substrates subtending these spectral fingerprints. Nonetheless, these correlates might provide an unique opportunity to infer potential mechanistic explanations, which at this point remain speculative and in attendance of further evidence.

We here hypothesize that complex fingerprints (i.e., multi-modal, with several power peaks) emerging in response to single pulse electrical stimulation could be generated by site-specific cross-frequency oscillatory mechanisms and reflect the regulatory role of local structures operating in the stimulated region. Recent physiological studies in non-human primates have revealed a hierarchical cortical regional organization with regards to the operational time-scales allowing (hence also limiting) intrinsic spiking fluctuations. To this regard, it has been shown that sensory areas exhibit shorter time-scales enabling a rapid detection of stimuli ^35^. In contrast, higher order regions such as pre-frontal areas operate with longer time-scales to integrate larger volumes of information, enabling more sophisticated and also time-consuming computations, such as for example in decision making^36^. In the human brain, electrophysiological studies seem to argue in favor of a regional organization with multiple time-scale rhythms ^37, 38^. Moreover, MEG recordings have shown that whereas some human cortical regions operate within well-defined frequency bands (i.e., finely tuned narrow-band spectra), other areas are less frequency selective and operate more broadband^39^. Supporting a non-hierarchical organization of neural activity, human brain regions do not necessarily follow a strict low-to-high order of activation time-scales and are able to operate simultaneously at multiple and differentiated frequency-bands. In this framework, we hypothesize that the level of complexity of site-specific spectral fingerprints reported in our study reflects the emerging properties of local systems, hence it could be considered a *proxy* to study by inference, the structural and functional (time-scales) organization of the sampled cerebral locations.

Additionally, complex spectral fingerprints could emerge from mixed contributions of locally generated ‘natural’ frequencies (reflecting the neurophysiological properties and local connectivity of regional oscillators) and rhythmic input from oscillators hosted in distant areas, connected with the stimulated region. Importantly, the role of the latter mechanism via signal reentry would increase in influence and spatial span as the cycles post-perturbation (i.e., in our case an electrical pulse onset) elapse, allowing more time for large-scale feedback and recurrent interactions. In agreement with this view, local influences should have a major role on spectral power changes during early post-stimulation cycles (1^st^ and 2^nd^ cycles), whereas network influences (presumably via re-entry mechanisms) might have most notorious bearing at longer time-intervals (corresponding to later frequency cycles), allowing the influence of local oscillatory processes by feedback projections.

Finally, the complexity of spectral fingerprints could also depend on the topology of the connectivity patterns sustained by the stimulated site with other brain regions and be particularly influenced in relay areas coping with a high volume and variety of inputs from widespread distant areas. In this vein, prior studies using resting state MEG activity in humans have revealed that similarity of spectral profiles across different brain areas might resemble resting-state cerebral networks such sensorimotor, frontal and visual networks, supporting that spectral fingerprints might show a hierarchical organization coherent with basic anatomical cortical subdivisions ^40^. In support of this idea, our data show that cortical sites with complex fingerprints (e.g. Angular Gyrus, Pre-cuneus and Middle Frontal cortices) are hosted in regions which have been identified as functional “hubs” of the default mode network (DMN) ^41^, and associated with high-level cognitive functions such as internal mentation, social working memory, autobiographical tasks, theory of mind, moral reasoning and episodic memory, among many others ^42^. In other words, high-level cognitive operations and processes need to rely on the integration of information through the different functional hubs of the DMN, hence operate at different time-scales and cope with rhythmic multiband activity.

In sum, we hypothesize that the morphology of the regional oscillatory fingerprints characterized in our study could allow us to better understand the likelihood of brain regions (as components of functional networks) to be synchronized at specific frequency bands to subtend specific behaviors. Nonetheless, the latter possibility calls for further evidence able to associate specific brain structural and functional connectivity properties across local and extended networks with the complexity of local spectral fingerprints and their cognitive contributions.

### General methodological considerations

We here benefited from the unique stimulation focality and signal-to-noise ratio of intracranial approaches to perturb and record rhythmic brain activity. Nonetheless, the use of iEEG data from epilepsy patients suffers some limitations. First, multielectrode implantation schemes and contact position differ greatly across individuals, hence a comprehensive map of ‘natural’ frequencies requires the integration of data from large cohorts of individually implanted patients. In our study, the combined implantation schemes of n=19 epilepsy patients achieved quite widespread anatomical coverage. Nonetheless due to the exquisite focality of intracranial stimulation and recording techniques (in the order of 5-10 mm), our current atlas of ‘natural’ frequencies remains sparse and needs to be further enriched by additional datasets, particularly from patients implanted in under-sampled non-temporal brain regions.

Second, to keep clinical mapping sessions within a reasonable duration and minimize the number of pulses and the total cumulated electrical charge undergone by patients, neurologists re-test the impact of electrical stimulation using identical parameters (i.e., same site, same pulse intensity and same burst frequency) at best, a few times for each pair of adjacent contacts. The scarce number of repetitions requires analyses at the single trial level and the averaging of datasets from the same or different patients, across stimulation intensities, and multi-electrode contacts hosted in different sites but laying within the same AAL regions (which in some cases can be very large). These approaches have been shown effective to cancel out spurious non-consistent spectral effects. Nonetheless, they also reduce the spatial resolution of the mapping effort and increase the risk of biases caused by high inter-individual variability, non-linear responses to increasing levels of electrical stimulation or architectural anisotropies within AAL atlas sites from which data end up clustered together.

Third, the lack of low-frequency activity (alpha and theta band) in our spectral fingerprints could be influenced by the Laplacian spatial derivative applied to recorded signals during pre-processing. This procedure removes signal components common across contacts within the same multielectrode. Accordingly, slow oscillatory components likely related to synchronization of larger neuronal population could be diminished and be difficult to detect. However, the Laplacian derivative is the method that minimizes inter-channel correlations in the iEEG time courses, necessary for atlasing purposes^43^. Other approaches commonly used to re-reference signal in similar intracranial EEG montages relay on the use of a common reference (intra- or extra-cephalic), removing *off-line* signals recorded by a specific contact from remaining recording contacts. This method enhances differences of recorded voltage, but is spatially unspecific and might blur accurate localization or eventually exaggerate irrelevant voltage differences. In the last years, a novel re-referencing method consisting in removing contact-wise the signal recorded in the closest white matter location has been probed ^23^. This approach might prove more accurate than the use of a common reference to localize voltage changes, but has also the risk of neglecting small voltage shifts and behave unequally across contacts, as the distance to the closest white matter is not always the same across all contacts.

Fourth and last, our time-frequency analyses required the suppression of “noisy” components associated to electrical stimulation artifact. We did so by removing 12 ms of iEEG data around stimulus onset and interpolating iEEG signals across this period with a third-degree spline. The methodological reliability of this method, initially developed to process concurrent TMS-scalp EEG recordings^14^ has been extended and validated for intracranial stimulation studies on “artificially” artifacted real and/or modeled EEG datasets, by showing that the oscillation power remains identical following artifact cleaning ^16^. Nonetheless, the removal of 12 ms of post-stimulus electrically artifacted data precludes reliable analyses of power increases at frequencies equal or higher than 80 Hz, since a complete cycle of real iEEG is eliminated by artifact cleaning procedures. Consequently, we can neither confirm nor rule out the presence of gamma components higher than 80 Hz in our spectral fingerprints.

### Concluding remarks and future directions

We here developed and tested a procedure based on direct electrical brain stimulation combined with intracranial EEG recordings of the human brain. We used such datasets to characterize and map the topographic distribution of local ‘natural’ rhythmic activity in the human brain. Our measurements could have likely capitalized on previously reported phase-resetting properties of focal single electrical perturbations, revealing the frequencies at which brain regions might most likely synchronize spontaneously.

From a fundamental research perspective, by characterizing the complexity of site-specific ‘natural’ spectral fingerprints, our study has generated testable hypothesis on the anatomical and neurophysiological properties of local oscillators in the human brain. Nonetheless, future intracranial datasets and analyses are needed to further enrich our understanding on why specific sets of local or distant brain regions might be more likely to get synchronized, why this might happen at specific frequency bands, and determine which sites share most similar or compatible spectral fingerprints, hence could be more likely to get coupled in phase or frequency. Moreover, by deepening our knowledge on how local oscillatory properties influence coding locally and across large-scale interregional systems, we will also be able to better understand the mechanism by which local and interregional synchrony enables specific cognitive processes and behaviors in the human brain.

From an applied technical perspective, invasive (e.g. deep brain stimulation and intracranial stimulation) and non-invasive (e.g. repetitive TMS and tACS) rhythmic brain stimulation have proven efficient at boosting ‘natural’ oscillations, particularly when tailored in phase and/or tuned to the ‘natural’ frequency at which a given area is most likely to rhythmically fluctuate. To this regard, accurate site-specific causal maps of local ‘natural’ oscillatory frequencies have the potential to enhance our understanding of current exploratory, diagnostic and therapeutic uses of invasive (e.g. therapeutic deep brain stimulation in Parkinson’s disease, dystonia or obsessive-compulsive disease or diagnostic intracranial stimulation in epilepsy) and non-invasive brain stimulation methods, and optimize future uses for the modulation of normal and pathological cognition.

## ACKNOWLEDGEMENTS

This article is dedicated to Lena Amengual Rocamonde, who came to this world in the middle of the production of this article. Authors thank Prof. V. Navarro, Dr. K. Lehongre, V. Lambrecq, Dr. S. Withmarsh, Dr. V. Frazzini and the CENIR EPIMICRO platform for conceptual, logistic an analytical assistance and for critical comments at different phases during the development of this project. Special thanks also for Dr. Mario Valderrama PhD, for allowing the use of custom-developed software to display electrode contact location data reported in this manuscript. We finally also thank Naturalia & Biologia Foundation for financial assistance for traveling and attendance to meetings. The activities of Dr. Valero-Cabré laboratory are supported by research grants IHU-A-ICM-Translationnel, Agence National de la Recherche (ANR), projet Générique “OSCILOSCOPUS”, the PHRC Regional “NEGLECT” and the PHRC Regional “STIM-SD” and eraNET-JTC-HBM CAUSALTOMICS. The contribution of Ms. Chloe Stengel is funded by a PhD fellowship of the École Doctorale ED3C at UPMC-Paris VI University. Dr. Julià Amengual was funded by FYSSEN Foundation.

